# Self-centering steady-state flows emerge in confined actomyosin networks

**DOI:** 10.64898/2026.09.24.754221

**Authors:** Jianguo Zhao, Charlie Duclut, Abhinav Singh, Rahil Golipour, An Pham, Behzad Golshaei, Chonglin Guan, Mingru Li, Ulrike Schulz, Rudolf Oldenbourg, Ivo F. Sbalzarini, Stephan W. Grill, James L. Harden, Frank Jülicher, Christoph F. Schmidt

## Abstract

The actin cytoskeleton drives shape changes and transport in cells and supports mechanical signal transmission. However, how cells control and use active cytoskeletal flows at the mesoscale is not well understood. We reconstituted an active cytoskeleton in water-in-oil emulsion droplets of *Xenopus laevis* egg extract and observed, above a critical droplet size, the emergence of a 3D radially convergent steady-state flow of polymeric actin, maintained by continuous actin turnover. The flow condensed lipid-rich cellular debris into a centered inclusion. Steady-state F-actin density and flow velocity profiles roughly collapse onto scale-invariant master curves. This behavior can be explained by a physical model representing the network as an isotropic active viscous fluid with a percolation threshold. The contracting network behaves as an active swimmer with complex internal dynamics that centers itself and the central inclusion inside the droplets without physical boundary attachment. Active contraction, crosslinking and polymerization dynamics in an actin network can thus generate cell-scale flow patterns that sense the confining geometry and external signals and exert forces that are likely sufficient to move and localize organelles in cells.

## MAIN TEXT

Individual eukaryotic cells as well as multicellular tissues display emergent mesoscale dynamics driven by molecular force generation. Such dynamics drive cell shape changes, organelle positioning within the cytoplasm, cell locomotion, cell division, as well as large-scale tissue movements in developing embryos (*1*). Collective dynamics driven by dispersed energy transduction, leading to force generation are a hallmark of active matter (*2*). In cells, emergent dynamics involve coordinated chemical and spatial fluxes of molecular players, which include both the force generators (molecular motors) and the structural filaments of the cytoskeleton. The actin cytoskeleton and non-muscle myosin II motors are the dominant force generators in many of these processes. Both are controlled by an array of enzymes such as actin crosslinkers, polymerizing and depolymerizing factors, and myosin activators or deactivators (*1*). A major challenge is to understand the connection between molecular interactions and emergent mesoscale collective dynamics. Cell-scale collective dynamics determine cell shape, for example for cells migrating on surfaces (*3*), but are also sensitive to boundary conditions and spatial confinements, imposed by an extracellular matrix or neighboring cells in tissues. These dynamics are thus part of the mechanosensory machinery of cells, generating and transmitting forces as well as reacting to forces. This type of mechanosensing is much less well understood than that performed by more static structures such as stress fibers and focal adhesions (*4*). Given the chemical and structural complexity of cells, it is practically impossible to capture all relevant molecular processes in *in vivo* experiments. There has also been limited success in reconstituting physiological processes in well-controlled systems with only a few components. For example, mixtures of purified and reconstituted microtubules and kinesin motors can form patterns such as asters or vortices (*5*), but cannot recapitulate mitosis. Purified and reconstituted filamentous actin with added myosin-II motors were shown to contract, a phenomenon originally dubbed “super-precipitation” before the contractile behaviors were understood (*6, 7*). But in this case the final states are rather static structures that are very different from continuously moving cytoskeletal structures in living cells.

An intermediate approach are experiments using cell fragments or cell extracts. Keratocyte fragments can show locomotion (*8*), and *Xenopus laevis* frog egg extracts have been extensively used to study mitotic and meiotic spindle formation and chromosome separation (*9, 10*). *Xenopus* extracts were shown to also produce collective myosin-driven actin flows (*11*), including cortical flows (*12, 13*), and radial bulk flows (*14–16*), similar to what is seen in various oocytes (*17–20*). It is intriguing to ask if it is possible to identify simple unifying principles explaining the characteristic features of the observed collective dynamics and tie them to a robust theoretical framework, with the final goal of connecting the underlying molecular processes to the mesoscopic dynamics.

We here constructed centro-symmetric contractile 3D F-actin flow patterns in water-in-oil emulsion droplets containing M-phase cytoplasmic extract of *Xenopus laevi*s eggs (Fig. 1A) (*21*). We studied F-actin velocity and density profiles and found scale-independent steady-state spatial patterns above a critical droplet size (Fig. S1). The dynamic patterns remained stable for hours until ATP likely ran out. Using tracer particles of different diameters, we showed that the background solvent remained quiescent, implying free-draining flow of F-actin towards the center of the droplet, establishing a stationary density gradient of F-actin. Lipid-rich cellular debris left in the extract was swept towards the droplet center and formed a condensed inclusion resembling a cell nucleus. Displacing the inclusion by magnetic beads showed that the centering remains stable to such perturbations.

**Fig. 1.**
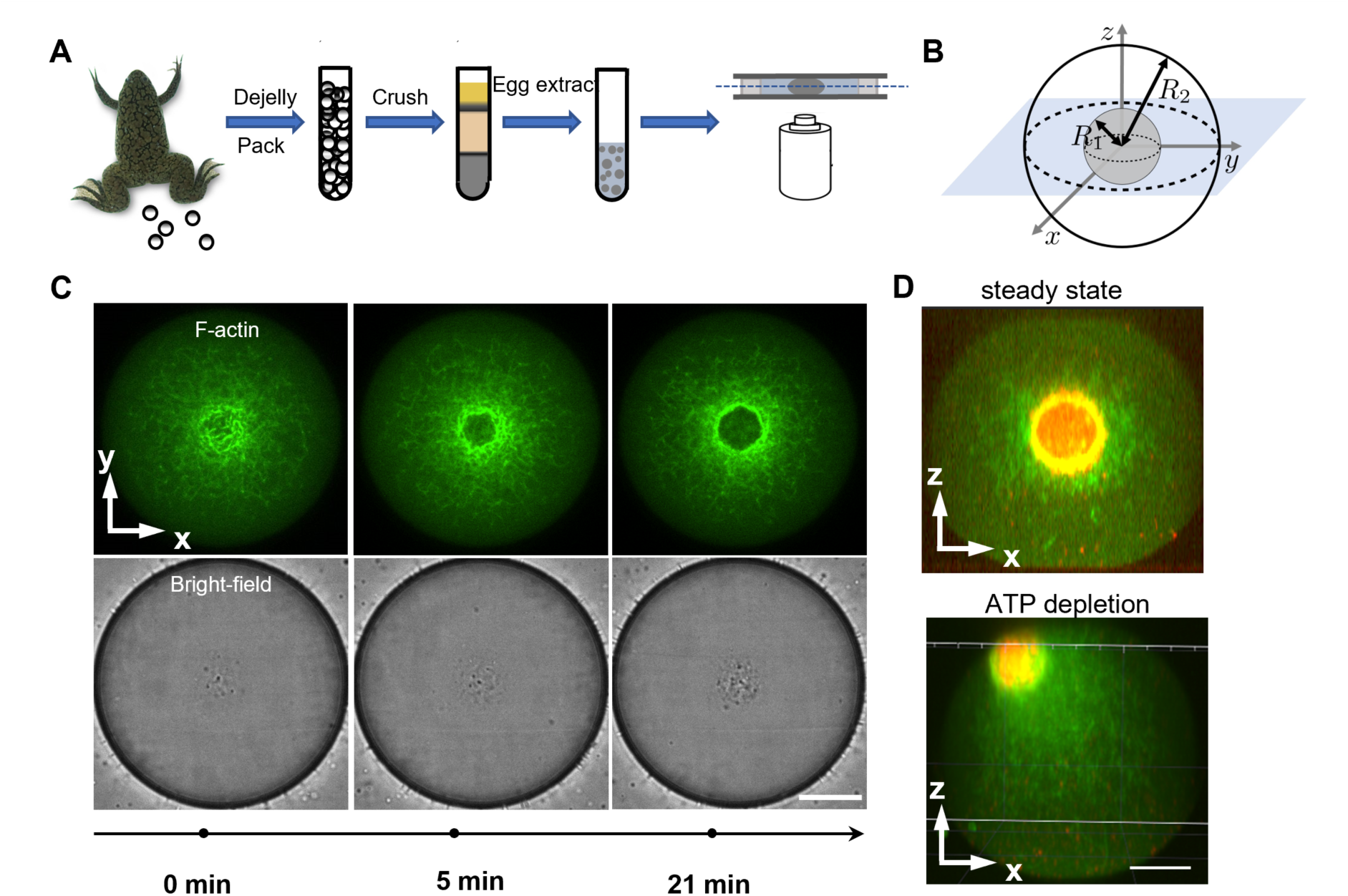
Self-organized steady-state F-actin flow centers inclusions in confined droplets. (**A**) Preparation of water-in-oil emulsion droplets containing *Xenopus laevis* egg extract. Cytoplasmic extract was encapsulated in mineral oil to form droplets confined between passivated glass slides separated by a 70-µm spacer. (**B**) Schematic of a confined droplet of radius *R*_2_ containing a lipid-rich inclusion of radius *R*_1_. The equatorial imaging plane is indicated in blue. (**C**) Time-lapse snapshots of equatorial cross-sections of the actin network (LifAact-GFP, green; top row) and corresponding bright-field images showing vesicular debris concentrated in the center (bottom row). The initially disordered actin network self-organized into a steady-state radially convergent flow that transported cellular debris toward the droplet center to form a dense spherical inclusion. (**D**) Vertical cross-section of a representative droplet from a confocal stack. The lipid-rich inclusion (Cy5, red) remained centered under steady-state F-actin flow (top). Upon ATP depletion, the inclusion drifted toward the droplet periphery before the flow ceased entirely (bottom). Scale bar in (**C**) and (**D**), 20 µm.

To understand the underlying dynamics of the network flow, we constructed a coarse-grained hydrodynamic model that describes the contracting acto-myosin network as a one-component active fluid (*15, 17, 22–26*). We show that this system partitions into two regions: a sparse actin network in the periphery of the droplets, behaving as a noncontractile viscous fluid, and an increasingly dense contractile network in the interior region of the droplets. The model quantitatively captures the measured F-actin density and velocity profiles and their scale-invariance, indicating that droplet radius is the dominant characteristic length scale of the flow patterns.

To understand the observed stable centering of the flow patterns, we suggest that one can consider a contracting acto-myosin network in a dynamic steady state as an active swimmer. The concept of active swimmers has been originally developed to explain the dynamics of self-propelled colloidal particles or microorganisms including bacteria and amoebae (*27, 28*), more or less rigid objects with a means of propulsion in a fluid. Here we consider the whole actomyosin network in the droplets as akin to a swimmer driven by its internal dynamics (polymerization, depolymerization, contraction) that swims towards the geometric center of the droplet and thereby stably centers itself in its confining boundary and also centers the inclusion without an elastic connection to the wall, merely driven by the supply of building materials (actin monomers) and the solvent flow generated when the swimmer goes off-center.

### Emergence of centro-symmetric contractile F-actin flux in emulsion droplets

*Xenopus laevis* egg extract was prepared following standard procedures (*21*) as described in Materials and Methods. To perform experiments, we prepared samples from frozen material by mixing extract with mineral oil and surfactant, followed by magnetic stirring to create emulsion droplets varying in diameter between a few to 100s of µm (Fig. 1A). The emulsion was filled into simple glass-slide/cover-glass sample chambers with ∼70 µm inner height, so that larger droplets were slightly flattened and immobilized between the top and bottom of the chamber (Fig. 1, A and B). Actin polymerization was triggered by raising the sample temperature to room temperature (21 ± 2 °C) after the sample was mounted for observation in a confocal fluorescence microscope. F-Actin was visualized by labeling with purified *LifeAct-GFP*. Microtubule polymerization was inhibited by adding 30 µM nocodazole. Within minutes after mounting in the microscope at room temperature, dynamic F-actin patterns emerged in the droplets, the structure of which varied depending on the droplet size (see SI, Fig. S1). For droplets <10 µm in radius, actin filaments mostly assembled into aggregates that spanned the droplet diameter and showed myosin-driven non-equilibrium fluctuations, but no collective flow (SI, movie S1). For droplets of 10 - 15 µm in radius, F-actin preferably contracted to a single cluster, that often ended up attached to the surface of the droplets (SI, movie S2). In droplets larger than 15 µm, actin filaments initially assembled into a crosslinked homogeneous network. Within minutes, a centro-symmetric contracting F-actin flow pattern emerged which evolved in time toward a dynamic steady state (Fig. 1C, SI, movie S3). The contracting network transported lipid-rich cellular debris present in the extract towards the center where it eventually formed a roughly spherical inclusion (superficially resembling a cell nucleus), which, in droplets prepared from the same batch of extract, scaled with droplet diameter, with some variability from preparation to preparation (SI, Fig. S2). The debris that was swept towards the center appeared to largely consist of micron-sized vesicles visible in phase-contrast microscopy (Fig. 1C, bottom row; SI, movie S3) which was confirmed by fluorescence microscopy using a lipid dye Cy5 (Fig. 1D, SI, movie S4). F-actin inside the inclusion gradually disappeared, while outside of the inclusion a density profile developed that decreased from a maximum near the edge of the inclusion towards the droplet periphery (Fig. 1C, top row; Fig. 2, A and C, SI, Movie S3). The F-actin density profile became stationary within ∼20 min and remained in a steady state in the presence of an ATP regeneration system added to the extract (Fig. 1D, SI, movie S4 and S5). The steady-state flow patterns typically broke down after ∼2 hours, presumably when ATP was beginning to run low, but before motility completely stopped. Typically, at that point, the position of the central inclusion became unstable and the inclusion migrated to the droplet periphery (SI, movie S5). This phenomenon is likely due to eventual connectivity percolation all the way to the droplet surface that might be caused by the increasing duty ratio of myosin at decreasing ATP concentrations (*29*), which would lead to increased F-actin cross-linking.

**Fig. 2.**
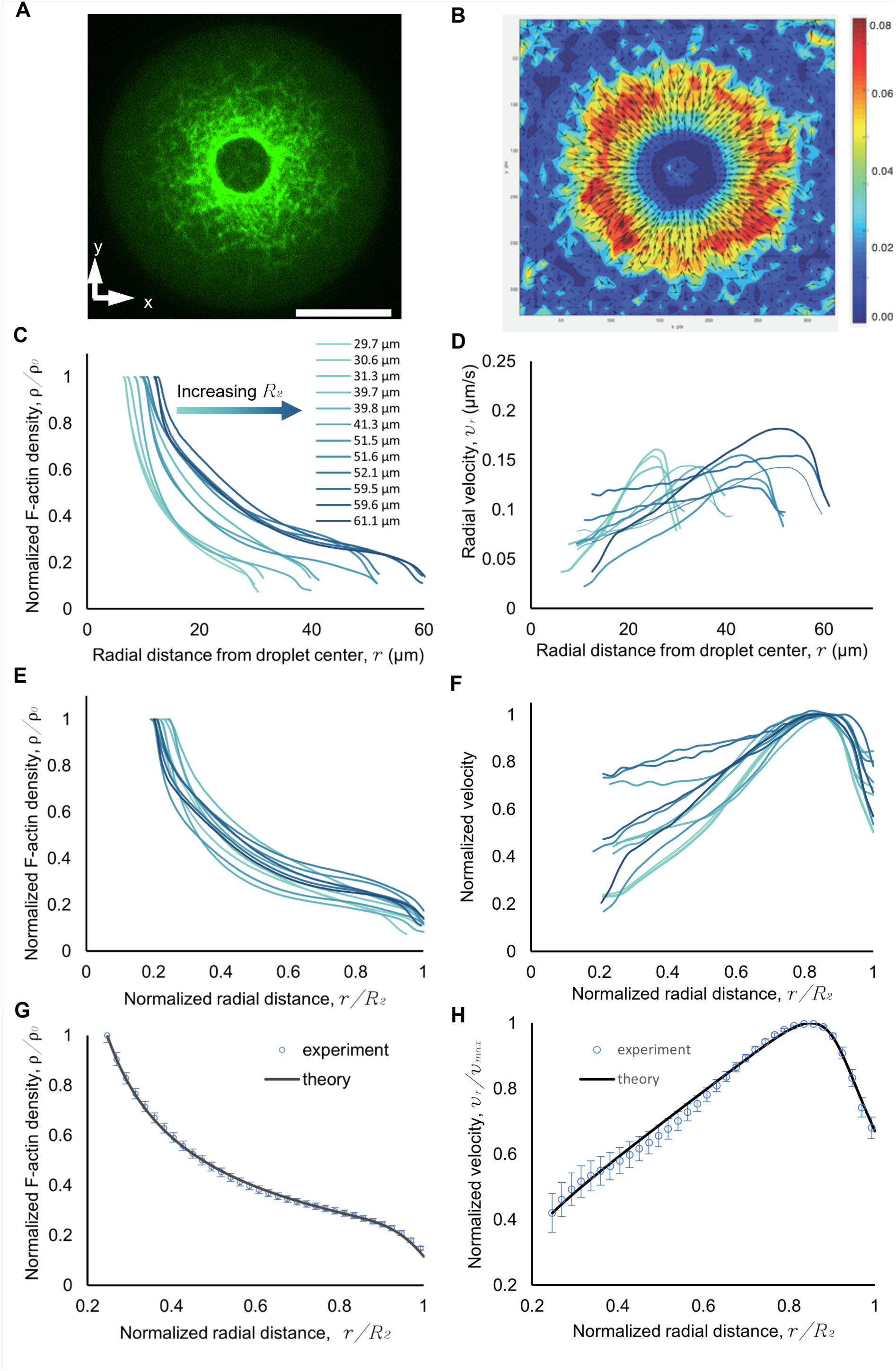
Quantitative characterization and hydrodynamic modeling of emergent F-actin flow. (**A**) Snapshot of F-actin distribution from fluorescence microscopy (LifeAct-GFP, green) during steady-state flow in a 60 µm emulsion droplet. Scale bar: 20 µm. (**B**) F-actin velocity field obtained by particle image velocimetry (PIV) and averaged over 200 s time windows. Color scale indicates velocity magnitude, arrows flow direction. (**C-D**) Azimuthally averaged F-actin density and radial velocity profiles measured in droplets of different sizes (**R_2_** = 29.7– 61.0 µm). (**E-F**) Corresponding profiles plotted as a function of radial distance normalized by droplet radius. Curves in (**C-F)** are color-coded by droplet radius (*R₂*), with darker shades corresponding to larger droplets. (**G-H**) Averaged normalized experimental profiles compared with model predictions (solid lines). Symbols represent averages from the 12 droplets shown in C - F. Simulation details and model parameters are provided in SI.

To find out if the radial flow pattern observed in the equatorial plane of the droplets was a convective vortical flow pattern, transporting F-actin back to the periphery in the z-direction or an isotropic 3D flow, we recorded z-stacks in a confocal microscope. Slices in vertical planes (Fig. 1D and SI, movie S4 and S6) show that the flow of the actin network was in fact radially convergent from all directions and not re-circulating. This observation implies that the steady-state inward flux of F-actin must be balanced by an outward flow of monomeric actin (G-actin), produced by filament depolymerization. This monomer flux was not visible in our fluorescence microscopy protocol. It is known that individual actin filaments undergo polymerization-depolymerization cycles on the minute timescale in the cytoskeleton of living cells and in cell extracts (*15, 16*).

If the network flow is close to spherically symmetric, the embedding fluid must be close to stationary because there is no sink of solvent in the center of the droplets and no source in the periphery. This means that the actomyosin network contracts through a stationary fluid background. Note that such free-draining flow is different from cytoplasmic streaming observed in oocytes (*19, 30, 31*), but is likely closely related to F-actin retrograde flow in locomoting cells (*32*). To confirm that the contracting network swept through a stationary fluid, we added 50 nm and 1 µm fluorescent beads to the extract and observed the transport of the beads driven by the contracting network (SI, movie S7 and S8). Some of the 50 nm beads ended up in the central inclusion, but many remained distributed in the whole droplet volume and diffused isotropically, largely uncoupled from the actin inward flux (SI, Fig. S3). Larger beads (1 µm) exhibited purely diffusive motion in the periphery of the droplets (in a ∼5 µm layer in ∼100 µm droplets), while they became strongly coupled to the radially convergent F-actin flux closer to the center of the droplets, eventually accumulating at the droplet center. This observation demonstrates (i) that the contracting F-actin network has a pore size smaller than 1 µm and (ii) that percolation of the polymerizing actin filaments into a network occurs at a distance of a few µm from the droplet surface. The latter implies that the contracting network is not physically connected to the water-oil interface and contracts with a free boundary condition. The transport of the larger particles in the contracting network provides a separation and selective transport mechanism that might be at work also in living cells when organelles are transported and localized.

### Absence of nematic order in the presence of rapid network turnover

Due to the fact that actin filaments and bundles of actin filaments are semiflexible and have a large aspect ratio, nematic ordering occurs in concentrated reconstituted actin solutions (≳2 mg/mL) (*33, 34*). Nematic order in active filament solutions has been shown to lead to large-scale flow patterns in microtubule solutions activated by kinesin motors (*35*) and also in concentrated actin solutions activated by myosin motors (*36, 37*). To test if there was nematic order in our contracting networks that might affect the F-actin contractile flux, we performed polarization microscopy (*38, 39*) that can report even small degrees of order since the optical anisotropy of actin filaments is large (*34*). We found no measurable orientational order in the bulk network, while there was strong edge birefringence on the water-oil interface at the outer edge of the droplets. (SI, Fig. S4).

### Radial density and velocity profiles scale with droplet size

In order to quantify the steady-state flow patterns and to construct an active fluid model describing the process, we first analyzed the F-actin network density profiles in horizontal equatorial cross-sections of the droplets at different time points after initiation of the contractile flow, using the fluorescence intensity of the LifeAct-GFP label which has a strong preference for binding filamentous actin over monomeric actin (*40*). As we found no persistent azimuthal density gradients, we azimuthally averaged the pixel intensities of fluorescent images in the equatorial plane to obtain the radial density profile at each time point (Fig. 2, A and C). We found a monotonically increasing F-actin density from the droplet periphery to the inclusion surface, indicating a transition from loosely organized actin bundles into a dense, crosslinked network (Fig. 2C). Notably, when plotted as a function of reduced radius, *r*/*R_2_*, where *R_2_* is the droplet radius, F-actin fluorescence intensity data for different size droplets at steady state approximately follow a master curve (Fig. 2E). Consistent with the observed motility of embedded 1 µm probe particles described above, the radial density profiles suggest a transition from a sparse, non-percolating peripheral network to a well-interconnected inner network closer to the droplet center.

Next, we analyzed the contraction velocity field in horizontal equatorial cross-sections of the droplets with particle image velocimetry, using the MATLAB implementation of OpenPIV (*41*), as described in Materials and Methods (Fig. 2B). We again assumed spherical symmetry and averaged the radial velocity components over all angles, thereby neglecting tangential velocity fluctuations (which we found to be less than the detection noise). The azimuthally-averaged velocity profiles approached zero at the surface of the inclusions and at the edge of the droplets as expected under the given boundary conditions. In the volume of the droplets, between inclusion and periphery, the velocity profile was non-monotonic with a maximum skew towards the periphery of the droplets (Fig. 2D). We found maximal contraction velocities between 0.11 and 0.18 µm/s, with an average of 0.15 µm/s, comparable to studies in similar reconstituted systems *in vitro* (*15*), and in starfish oocytes and *C. elegans* zygotes *in vivo* (17, 18). In contrast to these earlier results, however, we found non-monotonic velocity profiles. Interestingly, when the azimuthally-averaged radial velocity data is plotted as a function of reduced radius, *r*/*R_2_*, all data for different size droplets again collapse onto a master curve (Fig. 2F), consistent with the data collapse observed for the F-actin density profiles (Fig. 2E). These results show that the droplet size is the dominant length scale in the system.

### A hydrodynamic model for the contractile flow

In order to understand how a transient dynamic network made out of rapidly growing and shrinking filaments, crosslinked by accessory proteins and activated by contractile force dipoles can spontaneously generate the observed robust collective flow pattern, we developed a three-dimensional hydrodynamic continuum model whose steady states were compared to the experimental master curves (Fig. 2, G and H, Fig. 3). We describe the polymerized actin as a one-component active fluid with an F-actin density *ρ* and a velocity *v*. F-actin polymerizes and depolymerizes while its density is transported by the flow, such that it obeys the balance equation:

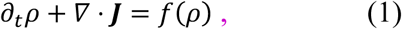

where ***J*** = *ρ**v*** is the F-actin flux and *f*(*ρ*) is an F-actin source term that describes the actin turnover, i.e. the balance between polymerization and depolymerization. At steady state, *f*(*ρ*) must be equal to the divergence of the F-actin flux, **∇** ⋅ ***J***, which can be computed from the experimental data (Fig. 3A). A steady-state radial inward flux of the F-actin network and a simultaneous outward flux of G-actin implies that there is net polymerization of actin in the periphery of the droplet, while depolymerization must dominate over polymerization towards the center, which is indeed observed (Fig. 3A). Having measured both density and velocity profiles, we can directly determine the F-actin flux ***J***(*r*) and calculate its divergence **∇** ⋅ ***J*** to quantify actin polymerization and depolymerization as a function of distance from the droplet center. We find a clear transition between a net polymerization zone near the droplet center and a net depolymerization zone near the periphery of the droplet. Furthermore, one can plot the flux divergence as a function of the local actin density, which shows that polymerization dominates (**∇** ⋅ ***J*** > 0) at low actin density and depolymerization dominates (**∇** ⋅ ***J*** < 0) at high actin density (Fig. 3A).

**Fig. 3.**
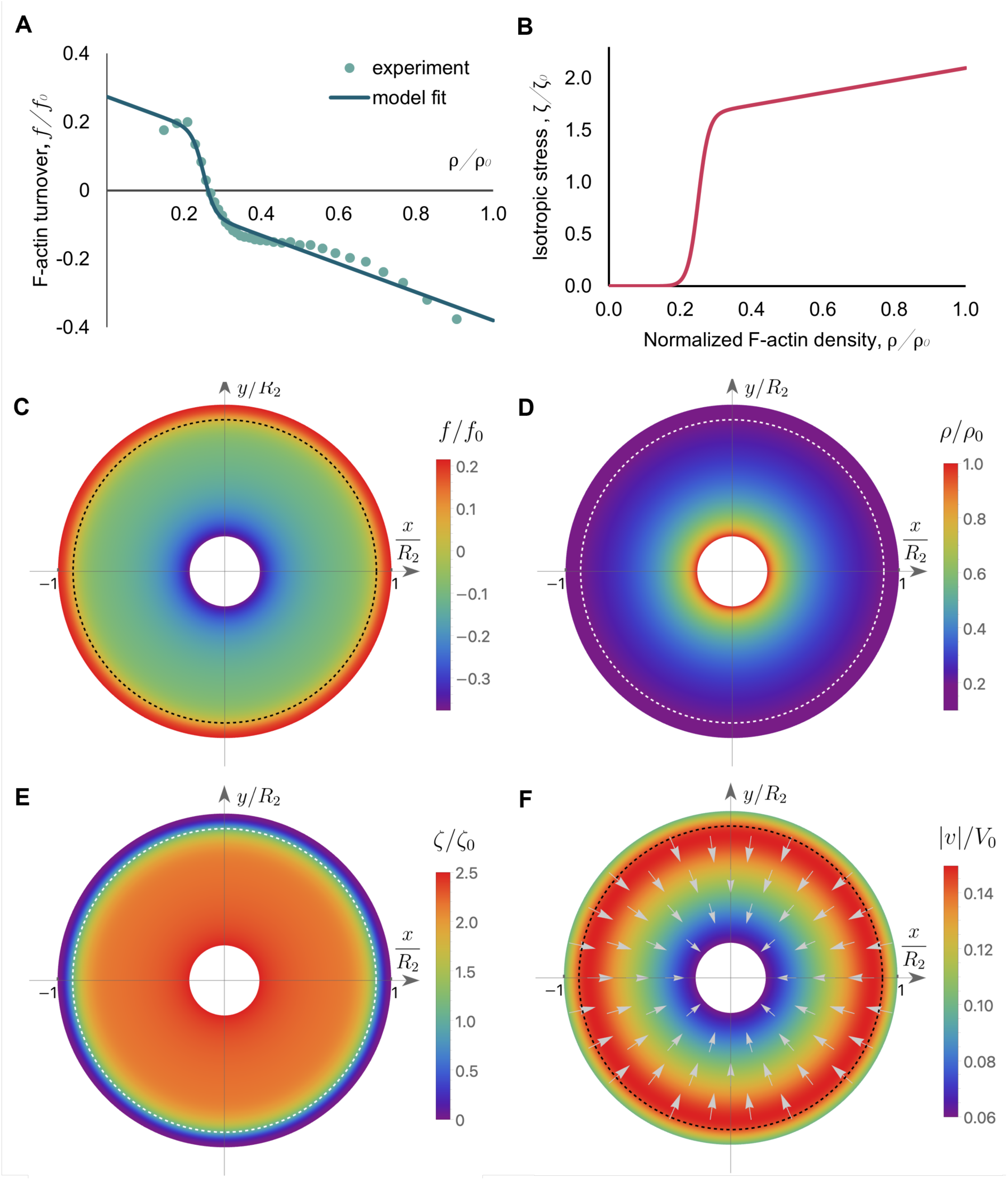
A 3D active-gel model reproduces the steady F-actin flow profiles through density-dependent turnover and stress. **(A)** Normalized F-actin turnover *f*/*f*_O_ as a function of normalized F-actin density *ρ*/*ρ*_O_. The experimental points are obtained by computing the steady-state F-actin flux divergence **∇** ⋅ (*ρv*) and are then fitted using Eq. (2) of the main text to obtain model parameters. **(B)** Normalized isotropic stress *ζ*/*ζ*_O_ as a function of normalized F-actin density *ρ*/*ρ*_O_. Active stress is generated only above a critical F-actin density threshold. (**C**) Normalized F-actin turnover profile along the equatorial (*xy*) plane. (**D**) Normalized F- actin density profile along the equatorial (*xy*) plane. (**E**) Normalized isotropic active stress profile along the equatorial (*xy*) plane. (**F**) Normalized F-actin velocity profile along the equatorial (*xy*) plane. In panels (**D-F**), the dashed line indicates the position *r_c_* at which the threshold density is reached (*ρ*(*r* = *r_c_*) = *ρ_c_*). Details of the numerical simulations and parameter values are given in SI.

To account for this transition between net polymerization and net depolymerization, we consider the following form for actin turnover:

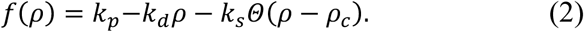

Below the threshold density *ρ_c_*, F-actin is polymerized from G-actin at constant rate *k_p_* (we neglect G-actin gradients), and depolymerization depends linearly on density with depolymerization rate *k_d_*. Above the threshold *ρ_c_*, a stress-induced depolymerization with rate *k_s_* leads to a net depolymerization. Near the threshold, the change is assumed to be sigmoidal and described by the nonlinear function *Θ*(*ρ*) = [1 + *tan*ℎ(*ρ*⁄*w*)]⁄2 with width *w*, which models a smooth transition between the two regimes. A fit (Fig. 3A) provides the threshold density *ρ_c_*/*ρ*_O_ ≃ 0.22 and the crossover width *w*/*ρ*_O_ ≃ 0.03 where *ρ*_O_ is the maximal F-actin density (See SI for details and fitted values of the rates).

Next, we formulated the forces acting on the network, i.e. the stresses in the continuum model. We constructed an active fluid model that accounts for the transition between a sparse non-contracting acto-myosin network in the droplet periphery and a contracting percolated network closer to the center. Force balance in the system reads as **∇** ⋅ ***σ*** = −*γ**v***, where ***v*** is the relative velocity between F-actin and solvent and *γ* is a coefficient describing the friction between the acto-myosin network and the fluid. The stress tensor ***σ*** appearing in the force balance is that of an isotropic active fluid:

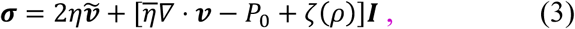

where ***ṽ*** is the traceless symmetric part of the actin velocity gradient, ***ṽ*** = (**∇** ⊗ ***v*** + (**∇** ⊗ ***v***)*^T^*)⁄2 − ***I*** (**∇** ⋅ ***v***)⁄3, in which ***I*** is the identity tensor in three dimensions, and *P*_0_ is a constant pressure. The shear and bulk viscosities *η* and *η* reflect the constant turnover of the F-actin network and can be estimated from the F-actin compression modulus *E* and its turnover time *τ* as *η* ≃ *η̅* ≃ *Eτ*. The relative importance of fluid friction and network viscosity is captured by the permeation length 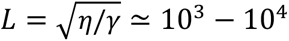, which is much larger than the system size. We thus consider *γ* = 0 in the following.

The active nature of the gel is captured by the isotropic active stress *ζ*(*ρ*). We postulate that below the percolation threshold, the network behaves as a non-contractile viscous fluid, while at larger density, the F-actin network is sufficiently dense to develop myosin-generated contractile stresses that drive actin flow. Based on these considerations, the isotropic active stress is written as:

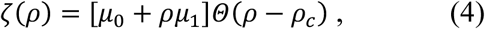

which captures the nonlinear transition between a sparse and passive regime (*ζ*(*ρ*) ≈ 0 at low density), and a dense and contractile regime (*ζ*(*ρ*) ≈ *μ*_0_ + *ρμ*_1_ is linear at high density, Fig. 3B). Note that we implicitly assume that the same physics – driven by a density threshold – underlies the nonlinear actin turnover rate of Eq. (2) and nonlinear active stress of Eq. (4), and we use the same sigmoidal function describing the threshold for both. This choice is validated below by comparison with the experiments. Similarly, a linear contractile regime at high density *ζ*(*ρ*) ≈ *μ*_0_ + *ρμ*_1_ is the simplest form that allows us to model the experiments accurately.

We then numerically solved the system of coupled nonlinear partial differential equations either considering a spherically-symmetric system (Figs. 2 and 3) or in full 3D (Fig. 4 and SI, movies S12-S14), using a numerical solver previously described (*25*). In both cases we used a fixed-velocity boundary condition from experiments, although a stress-free boundary condition can also be used with similar results (see SI for details of the numerical methods). The numerically obtained steady-state density and velocity profiles indicate that actin polymerization dominates over depolymerization at the periphery of the droplet, where the F-actin density is low (Fig. 3, C and D). When it reaches its threshold density (the corresponding position is indicated by a dashed circle in Fig. 3, C-F), the network exerts contractile stresses that create an inward F-actin flow (Fig. 3, E and F). The flow velocity is maximal close to this critical radius, where the gradient of active stresses is maximal (Fig. 3B), and then decays towards the center due to viscous dissipation and the reduced active stress gradient. Consistent with this nonuniform velocity profile, we note that the contraction rate (or velocity divergence) is non-monotonic along the radial position (Fig. S5). The F-actin flow results in increasing F-actin density as it accumulates towards the inclusion. Importantly, stress-induced depolymerization dominates (Fig. 3C) above the threshold density where internal stresses in the network are high. A stationary state can therefore be reached, when depolymerization exactly balances the F-actin flux transported by the steady-state flow.

**Fig. 4.**
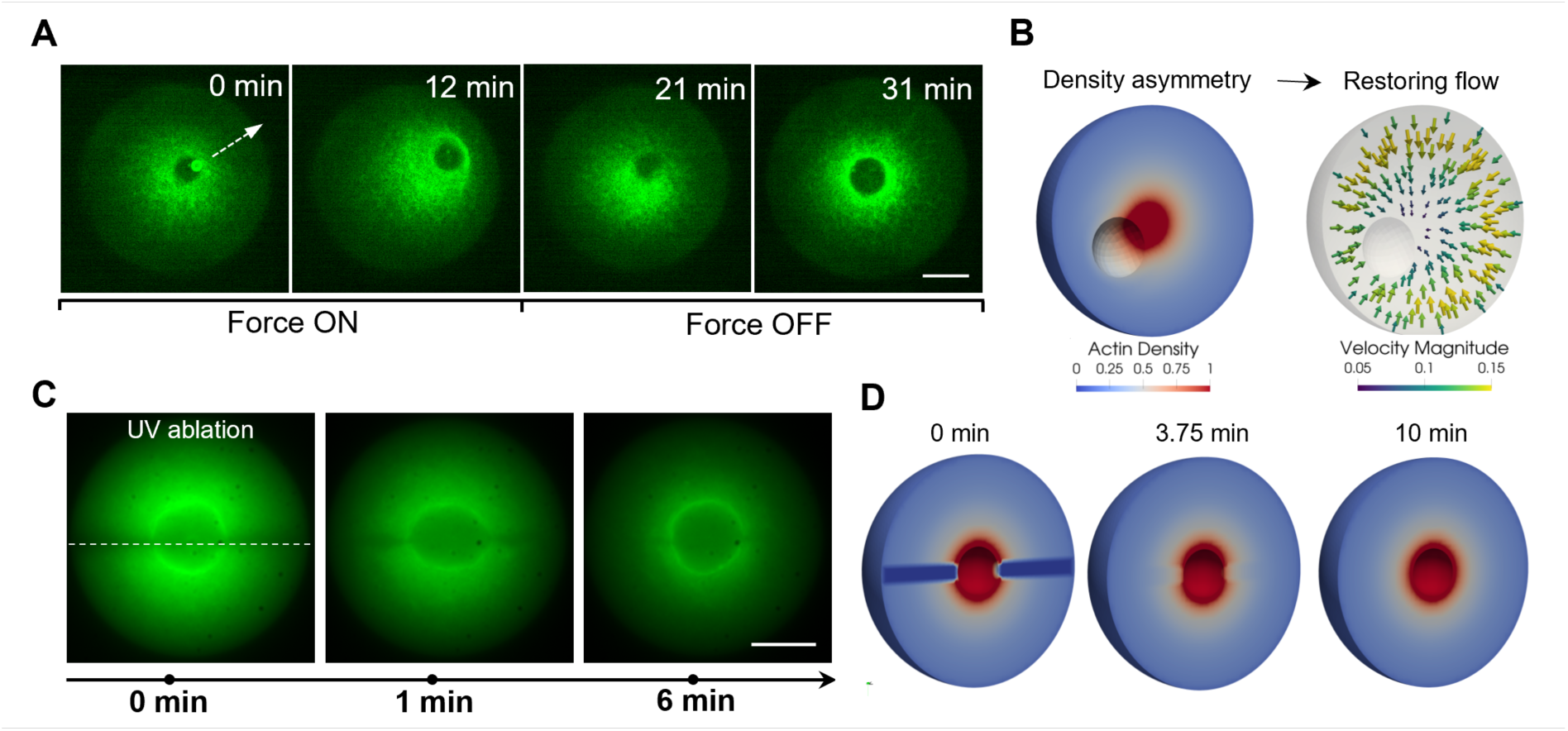
Recovery of the actin flow pattern after perturbations. (**A**) Displacement of the inclusion by an embedded magnetic bead (arrow) attracted by an external magnetic field (∼300 pN force). During displacement, the actin network (LifeAct-GFP, green) becomes asymmetric. After the magnetic field is turned off, flow symmetry and inclusion centering are restored in the absence of physical attachment to the droplet boundary. Scale bar, 20 µm. (**B**) Three-dimensional numerical solution of the steady-state F-actin density profile (left) and velocity field (right) around an off-centered inclusion. The inclusion is treated as a fixed and rigid object. The asymmetric density profile generates inward restoring flows that drive the inclusion toward the droplet center. (**C**) Recovery from UV laser ablation along the droplet diameter. UV ablation transiently disrupts the actomyosin network in the bulk and at the inclusion surface. Immediately after ablation, the inclusion deforms and flow symmetry is broken. As the network reconstitutes, the centrosymmetric configuration and spherical inclusion shape are progressively restored. Scale bar, 20 µm. (**D**) Three-dimensional simulation snapshots of F-actin density (color-coded background) following ablation, showing recovery toward the steady-state density profile.

Finally, we fitted the calculated actin velocity profile to the experimental data to determine the model parameters *η̅*, *η*, *μ*_0_, *μ*_1_ (see SI for details regarding the number of parameters and their fitting procedure). The results (Fig. 2, G and H) show an excellent agreement of the shapes of the model curves with the experimental data, capturing accurately the non-monotonic velocity profile. We emphasize that the nonlinear transition between a sparse and a dense regime is essential to obtain accurate fits of both the velocity and density profiles.

### Mechanical perturbations reveal robust self-centering

We found that the self-organized centrosymmetric flow patterns generated by the contractile network were resilient to various perturbations, and the insight gained from the theoretical model allowed us to rationalize the mechanisms underlying this robustness.

To probe the robustness of the centering mechanism and the dynamics of relaxation back to steady state, we embedded 4.6 µm diameter super-paramagnetic beads in the extract, which got swept into the inclusion by the contracting network. Once the system had reached steady state, we turned on an external magnetic field gradient in brief pulses, using an electromagnet with a ferromagnetic core ending in a sharp tip, placed near the edge of a droplet. The calibrated force on the bead (maximally ∼300 pN) moved the inclusions towards the droplet periphery at a slow speed of 1.3 µm/min on average (Fig. 4A, SI, Fig. S6, movie S9). The F-actin distribution became asymmetric during the displacement of the inclusion, with initially the whole actin network getting shifted towards the droplet periphery and subsequently a higher actin density building up towards the center of the droplet. After turning off the field, the inclusion relaxed back to the droplet center roughly exponentially with a relaxation time on the order of 10 min, even when the displacement was as large as about half the droplet radius (Fig. 4A, SI, Fig. S6), while the network recovered its centrosymmetric actin density distribution. This procedure could be repeated several times during the lifetime of the steady-state flow pattern (Fig. 4A, SI, Fig. S6). The asymmetry in the F-actin distribution is also observed in a 3D numerical solution of the model where the inclusion is held off-centered, with an enrichment towards the center of the droplet (Fig. 4B, SI, movie S10). This density asymmetry produces an asymmetric active contractile stress profile (which can be intuited as a negative pressure profile) that drives an F-actin flow that pushes the inclusion back to the center of the droplet. The robustness of the radially-symmetric solution was further checked in our 3D simulations, where we checked that asymmetric initial conditions still lead to the radially-symmetric steady-state solution (SI, movie S11).

We found that this self-organized structure was also resilient to a partial destruction of the actin network by cutting regions of the network using a UV-laser. Performing a cut straddling the inclusion and destroying the F-actin network locally to the left and right of the inclusion led to a rapid expansion of the inclusion along the direction of the cut (Fig. 4C, SI, Fig. S7, movie S12 and S13) followed by re-polymerization of the network and subsequent recovery of the spherical shape of the inclusion. The expansion along the direction of the cut is understood as a consequence of the weakening of the contractile force exerted by the actin shell in the ablated region. The recovery of the network in the bulk after ablation was reproduced by our 3D active-fluid numerical solution (Fig. 4D, SI, movie S14). The numerical solution, which describes the inclusion as a nondeformable sphere, show that the inward F-actin flow velocity is reduced where the ablation is performed (Fig. 4D), and then recovers to its steady-state value as the bulk actin network repolymerizes. The UV-ablation experiments thus revealed that the inclusion gets compacted into a condensed spherical mass due to the accumulation of actin into a contractile shell at its surface.

Together, these results demonstrate that the self-centering capability of the steady-state contracting actin network flow is extraordinarily stable and resilient to external perturbations. It is remarkable that the percolated inner network, which is separated from the droplet boundary by a non-percolated layer, still senses the boundary and develops stresses that keep it robustly centered in the droplet.

Dynamic cytoskeletal networks reconstituted in emulsion droplets from *Xenopus laevis* egg extract are known to display robust large-scale dynamic flow patterns that resemble processes observed in cells (*11, 14–16, 23, 24*). Observed flow velocities are comparable to actin retrograde flow velocities observed in surface adherent cells (approximately 1.2-15 µm/min) (*32, 42, 43*), where, just like in the droplets, a contracting actin cortex moves through an approximately stationary background cytoplasm. Chromosome transport in starfish oocytes was also found to be orchestrated by a contractile actin network (*18*), creating a linear velocity profile reaching contractile velocities of ∼6 µm/min at a distance of ∼70 µm from the animal pole. Interestingly, cortical flow in *C. elegans* zygotes, which is also driven by a dynamic, contracting acto-myosin network, exhibits maximal velocities of ∼6 µm/min as well (*17*). In this case one would, however, not expect permeating flow of the thin cortical actin layer.

Reported F-actin flow patterns in emulsion droplets include stationary contractile flow in spherical (*14, 15, 23*), cylindrical (*16*) or elongated (*11, 24*) droplets, as well as pulsatile concentric flow in large droplets (*24*). It was shown that flow patterns can be modified by manipulating relevant molecular components including motors, crosslinkers, or capping proteins (*15, 16, 24*). All these results suggest that network contraction due to myosin motors acting on crosslinked F-actin networks generates the inward flux, and that the continuous depolymerization of F-actin to G-actin and outward diffusive flux of G-actin maintains the stable steady-state dynamics. Radially inward network flow was observed to concentrate larger material remaining in the coarse extracts into (metastable) spherical aggregates. Aggregates recenter after perturbations (*23*), but move spontaneously to the periphery in smaller droplets and with time also in larger droplets (*14, 23*). Several hydrodynamic models have been proposed to explain both the flow patterns and the centering of the aggregates (*15, 16, 23, 26*).

We confirm here that contractile stress in the actin networks in conjunction with actin turnover generate steady centro-symmetric flow patterns once there is connectivity percolation, provided that the droplets are large enough and that ATP concentrations are high enough. The occurrence of these flows was robust and independent of the particular batch of *Xenopus laevis* extract. In addition, we could observe the gradual formation of the inclusions that at first still contained F-actin which then depolymerized while incoming F-actin got blocked once the inclusions were sufficiently compacted. Further compaction was driven by a contractile shell of acto-myosin, which was evident when disrupting the shell by local UV depletion of F-actin. In small droplets (<10 µm in radius) fluctuating F-actin bundles formed without establishing large-scale flow, which was not reported before. We propose that the reason for this phenomenon is that the fluidization of the contracting network at long time scales by actin depolymerization (necessary to create the steady state flow) is scale dependent such that flow does not emerge when the droplet diameter is comparable to the size of transient actin clusters.

The fact that we find no evidence of nematic ordering (Fig. 1 and S4) is in striking contrast to strong local alignment found in *in vitro* reconstituted dense systems of molecular motors and stabilized microtubules and actin filaments (*35–37*) and flow-aligned cortical actin *in vivo* (*44*). The rapid turnover of filaments and their short average length might lead to a situation where isotropic myosin-generated forces dominate over interactions favoring nematic ordering. Tracking of embedded small and large tracer beads showed (i) that the background solvent was largely stationary as expected in a centrosymmetric flow pattern, and (ii) that the actin network became so dilute near the surface of the droplets that even the 1 µm beads were not transported inwards anymore. F-actin was still flowing towards the center in that region, but with velocity decreasing towards the periphery, suggesting a lack of percolation and contractility. This is confirmed by our theory that fits the observed velocity profile when a transition from non-contracting to contracting network is implemented. Importantly this implies a stress-free boundary condition around the contracting network, i.e. no force transmission to the droplet surface.

To explain the stable centering of the flow patterns, several models have been proposed in the literature, including (i) a hydrodynamic mechanism based on Darcy friction between the contracting network and the surrounding cytoplasm (*23*) and (ii) a “tug-of-war” mechanism based on the dynamic competition between inward contractility and boundary-associated attachments (*16*). Active diffusion–based models have invoked stochastic motor-driven fluctuations that generate effective pressure gradients without persistent directional flow (*30*). More recently, passive mechanical properties of the cytoplasm have also been proposed to stabilize the mitotic spindle once central positioning has been achieved (*45, 46*).

Here, we suggest alternatively that the stable centering of both, the whole network and the inclusions in our emulsion droplets should be regarded as two coupled, but separate phenomena. The centering of the inclusion is possible because of a system-spanning percolated contracting network, which is able to generate the stress gradients that maintain the inclusion centered, as we demonstrate in our 3D one-fluid model. The whole network senses the boundary due to the cut-off of G-actin building block supply at the droplet surface. When the network is decentered, stress asymmetry in the network will also create hydrodynamic solvent flow, pushing against the droplet surface and recentering the network. Solvent flow is not included in our active fluid model, but momentum conservation, of course, demands that, during recentering, momentum is transferred to the droplet surface by hydrodynamics. Eventual connectivity percolation all the way to the droplet surface, leading to migration of the network as well as the inclusion to the droplet surface, is likely caused by the diverging duty ratio of the non-muscle myosin motors with decreasing ATP concentration, approaching the bound rigor state (*47, 48*) which increases the effective crosslink density in the network.

The 3D hydrodynamic active one-fluid model we developed to describe the observed flow patterns is as simple as possible while still reproducing the observed network dynamics. It treats the actin network as an isotropic Maxwell fluid in the long-time limit where viscous dominates over elastic response. This viscosity, importantly, stems not from drag against solvent (*23*), but from the relaxation of transient elastic stresses in the network by F-actin and crosslink turnover. Locally isotropic contractile stress is generated by myosin activity that is proportional to actin density. Active stress gradients manifest as a negative pressure in the network that drives the centrosymmetric flow.

Previous models (*15, 24*) focused on the linear velocity profile (*v* ≈ *a* + *br*) seen close to the central inclusion which corresponds to a constant contraction rate (or velocity divergence), requiring the *ad hoc* assumption that the effective F-actin viscosity and the active stress have exactly the same density dependence. In contrast, we show experimentally that in our system the velocity profile is nonmonotonic (Fig. 2H), and the contraction rate is not constant (SI, Fig. S5). In the model we therefore introduced a threshold F-actin density at which contractile stresses set in. The nonlinearity introduced by this threshold is key to obtain a nonmonotonic velocity profile in the droplet periphery while still predicting a linear velocity profile close to the inclusion (Fig. 3). Our model displays universal steady-state profiles dependent on the reduced radial distance *r*/*R*_2_and thus confirms that the droplet radius is the only relevant length scale. This is consistent with the collapse of the density and velocity profiles after scaling with droplet diameter. It will be interesting to assess if the key nonlinearity introduced here can also describe the reported actin waves and spirals in reconstituted actin systems in very large droplets (*24*).

The full 3D numerical solutions that we used to solve our model extend existing theoretical work (*15, 16, 24*) that had focused on spherically-symmetric solutions or 2D geometries. Our approach allowed us to probe nontrivial geometries – including the recovery of the F-actin network after UV ablation (Fig. 4D and SI, movie S14), and the F-actin profile around a decentered inclusion (Fig. 4A, SI, movie S10) – and to explore the origin of the centering force and the robustness of the spherically-symmetric steady-state, (Fig. 4B and SI, movie S11).

We suggest that one can view the system as an active swimmer, which swims back to the center after perturbation. In contrast to colloidal active swimmers, this system is a distributed, continuously remodeling active network that is defined by its internally generated stress and flow fields rather than by a fixed internal geometry. It senses boundaries and centers itself and its internal flow pattern in the confining droplets without touching the walls, but rather feels the walls through the availability of its constituents (G-actin etc.). In other words, the centered configuration represents a dynamical attractor of the confined active gel. When the structure is dragged from the droplet center, and confinement becomes asymmetric, the F-actin-depleted region left behind fills with new F-actin which leads to an imbalance in active stresses arising from coupled gradients in filament connectivity, contractility, and turnover, pulling the inclusion back to the center, while recentering the whole network due to solvent flow generated by the asymmetry of the network.

We conclude that the contracting network forming in the extracts is a fascinating active viscoelastic material, showing the mechanical properties of a viscoelastic Maxwell fluid, appearing viscous on time scales longer than the actin and crosslinker turnover times (minutes), while maintaining long-range contractility gradients and stress profiles driving network flow. It is intriguing to regard the contracting networks as unconventional active swimmers that adapt their internal dynamics to external boundaries which then leads to hydrodynamic forces propagated through the solvent when internal stresses are unbalanced. In the simple geometry of spherical emulsion droplets, the flow patterns are centro-symmetric. In the more complex geometry of cells in tissues, the same general mechanism might be relevant for the transport, localization and ordering of organelles, including the nucleus. The sensitivity of the internal dynamics to potentially far away boundary conditions would also provide a mechanism for cells to sense and transmit external forces, generating an unconventional path to mechanosensing, independent of specialized anchoring structures.

## MATERIALS AND METHODS

### LifeAct-GFP2 his_6_ purification

#### Cloning of LifeAct–GFP2–His_6_

The cDNA encoding LifeAct–GFP2 was initially amplified by PCR from the pCAG-LifeAct-tagGFP2 plasmid (Ibidi). The primers NdeI-LifeAct-for (5′- GGCCCATATGGGTGTCGCAGATTTGATC-3′) and XhoI-GFP2-rev (5′-CGGCCTCGAGCCTGTACAGCTCGTCCAT-3′) were used to generate a LifeAct–GFP2 fragment flanked by 5′ NdeI and 3′ XhoI restriction sites. The PCR product and the pET-24b(+) vector were digested with NdeI and XhoI and ligated to yield a construct encoding resulting in an in-frame LifeAct–GFP2–His_6_ construct under control of the T7 promoter.

#### Bacterial expression

The pET-24b-LifeAct–GFP2–His_6_ plasmid was transformed into *E. coli* BL21-Gold, and transformants were selected on LB agar plates containing 50 µg/mL kanamycin. For protein production, a single colony was used to inoculate 20 mL LB medium supplemented with 50 µg/mL kanamycin as a preculture. 1000 mL LB–kanamycin was inoculated with the preculture to an OD_600_ of 0.1 and grown at 37°C until an OD_600_ of approximately 0.6 was reached, at which the temperature was lowered to 22°C and expression was induced with 200 µM IPTG when OD_600_ reached 0.9. Induced cultures were incubated at 22 °C overnight (16 h) to favor soluble expression. Cells were harvested by centrifugation (4,600 × g, 25 min, 10 °C), and pellets were frozen and stored in liquid nitrogen prior to lysis.

#### Lysis and Ni2^+^-affinity purification

Cell pellets were resuspended in 43 mL resuspension/lysis buffer (50 mM Tris–Cl pH 8.0, 250 mM NaCl, 10 mM β-mercaptoethanol) supplemented with EDTA-free protease inhibitor (Sigma), 0.2 mg/mL lysozyme, and 25 µg/mL DNase I. After thawing on ice, cells were lysed by probe sonication (3-time 10s pulses with 1 min cooling intervals on ice), sheared by passing 3 times through a G21 syringe cannula, and insoluble material was removed by centrifugation (20,000 × g, 20 min, 4 °C). The clarified supernatant was incubated with 2 mL pre-equilibrated Ni^2+^-NTA agarose resin slurry in resuspension buffer to allow binding of the His^6^-tagged LifeAct–GFP2 fusion. After 30 min binding at 10°C on an end-over-end shaker, the beads were washed three times with 10 mL wash buffer each (50 mM Tris–Cl pH 7.4, 250 mM NaCl, 10 mM β-mercaptoethanol, 10 mM imidazole) to reduce nonspecific interactions. Bound protein was eluted with elution buffer (wash buffer with 300 mM imidazole added) in nine 1 mL fractions.

#### Further processing and characterization

Protein purity was assessed by SDS–PAGE Bis/Tris gel followed by Coomassie Brilliant Blue staining, and protein concentration was determined by absorbance at 280 nm using the calculated molar extinction coefficient of GFP. Fractions 1-8 were pooled, supplemented with 10% glycerol, and snap-frozen in 100 µL aliquots in liquid nitrogen, and stored at −80 °C.

### Xenopus egg extract preparation

Egg extracts were prepared as described in Ref. (*21*) with minor modifications. Briefly, freshly laid *Xenopus laevis* eggs were rinsed with MMR buffer containing 5 mM HEPES, 0.1 mM EDTA, 100 mM NaCl, 2 mM KCl, 1 mM MgCl₂, 2 mM CaCl₂, pH 7.8, and dejellied using 2% (w/v) L-cysteine prepared in XB buffer containing100 mM KCl, 1 mM MgCl₂, 0.1 mM CaCl₂. The eggs were then washed with CSF-XB buffer containing 100 mM KCl, 2 mM MgCl₂, 0.1 mM CaCl₂, 50 mM sucrose, 10 mM HEPES, 5 mM EGTA, supplemented with protease inhibitors of leupeptin, pepstatin A, and chymostatin at 10 µg/mL each. The eggs were packed by centrifugation at 1,000 rpm for 15 s in a clinical centrifuge and subsequently crushed by centrifugation at 12,000 rpm for 15 min at 4 °C. The cytoplasmic layer was collected and supplemented with an energy mix containing 9 mM creatine phosphate, 1 mM ATP, and 1 mM MgCl₂, together with 50 mM sucrose and protease inhibitors at concentrations indicated above, and recombinant His₆-tagged LifeAct–GFP2 at a final concentration of 0.064 mg/mL. The extract was aliquoted, flash-frozen in liquid nitrogen, and stored at −80 °C until use.

### Magnetic bead preparation

The superparamagnetic beads (4.5 µm diameter, tosylactivated surface; Dynabeads M-450 Tosylactivated, Thermo Fisher Scientific) were covalently functionalized with concanavalin A (ConA), which binds carbohydrate residues on membrane-associated proteins enriched within the droplet inclusions. Briefly, 200 µg of Alexa Fluor™ 488–conjugated ConA was incubated with 1 mL of bead suspension (4 × 10⁸ beads) in 0.1 M sodium borate buffer for 24 h at room temperature under gentle tilting and rotation. The beads were then rinsed twice for 5 min at 4 °C with Ca²⁺/Mg²⁺-free PBS containing 0.1% BSA and 2 mM EDTA (pH 7.4). Residual tosyl groups were blocked by incubation in 0.2 M Tris containing 0.1% BSA (pH 8.5) for 4 h at 37 °C. The functional beads were subsequently washed using the same PBS/BSA/EDTA buffer and stored at 4 °C until use. For magnetic force perturbation experiments, 0.5 µL of the bead suspension was mixed with 9.5 µL of egg extract.

### Glass passivation

Glass slides and coverslips were sonicated for 10 min in 1 M KOH solution prepared in 25: 75 (v/v) mixture of water and ethanol, followed by two of of 10-min sonication steps in Milli-Q water. After drying overnight at room temperature, the glass surfaces were passivated by vapor-phase silanization with dichlorodimethylsilane (85126, Sigma-Aldrich, Germany) for 2 h in a desiccator. The silanized slides and coverslips were then rinsed with heptane (34873, Sigma-Aldrich) and sonicated twice in Milli-Q water for 10 min each. The passivated glass was then dried at room temperature and stored at 4 °C for up to several months before use.

### Emulsion preparation

Emulsions were prepared as described in ref. (*49*) with minor modifications. Briefly, 10 µL of egg extract was added to 1 mL of degassed mineral oil containing 4% (w/v) cetyl PEG/PPG-10/1 dimethicone (ABIL EM 90, Evonik Nutrition & Care GmbH) and emulsified by magnetic stirring for 30 s. Approximately 20 µL of the resulting emulsion was introduced between a glass slide and coverslip, separated by 58-μm-thick double-sided adhesive tape (3M). The chamber was sealed with VALAP, a 1:1:1 mixture of Vaseline, paraffin, and lanolin. This procedure generated spherical extract droplets, providing a confined, cell-sized geometry and facilitated optical imaging of the cytoplasmic interior.

### Confocal Microscopy

Time-lapse imaging of the actin network was performed using an Andor Dragonfly spinning-disk confocal microscope at room temperature, equipped with a 63× oil immersion objective (NA 1.47, Leica) and an Andor Zyla PLUS 4.2 megapixel sCMOS camera. Fluorescence excitation was provided by a 488 nm laser, and emission was collected using a 525 nm emission filter to visualize LifeAct–GFP2. Fluorescence excitation was provided by a 488 nm laser, and emission was collected using a 525 nm emission filter to visualize LifeAct–GFP2.

### Polarized light microscopy

Birefringence imaging was performed using an LC-PolScope (*38*) equipped with a liquid crystal–based universal compensator to modulate illumination polarization. Images acquired at multiple compensator settings were used to compute pixel-wise retardance and slow-axis orientation, providing quantitative measures of molecular order within the specimen (*39*). Retardance and slow-axis orientation maps were computed from polarization-resolved image stacks using the OpenPolScope system with ImageJ/Micro-Manager plugins (https://openpolscope.org).

### Magnetic tweezers

The magnetic tweezers setup consists of a solenoid coil and a cylindrical core made of highly magnetically permeable steel. The solenoid was 50 mm long and 20 mm in diameter, and was formed by 1000 turns of 24 AWG copper wire wound around a brass frame. The carbon steel core (5 mm in diameter and 150 mm in length) was enclosed by the solenoid. One end of the core was tapered to a sharp tip with a radius of less than 10 µm. This sharp geometry concentrated the magnetic field gradient near the tip, thereby enhancing the attractive force exerted on the paramagnetic beads.

The magnetic tweezers assembly was mounted on a Leica three-axis micromanipulator, enabling manual adjustment of the probe position and working distance before each experiment. A custom MATLAB script was used to generate voltage signals and interface with external devices through a data acquisition card (NI USB-6001, National Instruments). The output signals were amplified using a bipolar operational power supply (Kepco Power Supplies) before being applied to the solenoid coil to generate the magnetic field.

Magnetic forces were calibrated by tracking the motion of paramagnetic beads suspended in a silicone oil of known viscosity. As the particle moved toward the magnetic tip, the magnetic force was estimated from the Stokes drag force. The drag is given by

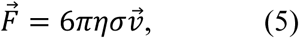

where *F⃗* is the force vector directed toward the tip; *η* is viscosity of silicon oil; *σ* is the bead radius, and *v⃗* is the particle velocity obtained from particle trajectories acquired by time-lapse imaging. The measured force magnitude was plotted as a function of the bead-to-tip distance and fitted with an exponential function

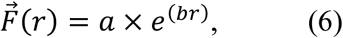

where *a* and *b* are fitting coefficients. The resulting calibration curve was used to estimate magnetic forces at the bead-to-tip distance extending beyond the microscope field of view (Fig. S8).

### UV laser ablation

UV laser ablation was performed using a diode-pumped solid-state laser (355 nm, approximately 0.84 µJ per pulse) controlled by a UGA-42 Caliburn scanning unit (Rapp Opto Electronic, Germany). The laser ablation system was integrated with a Zeiss Axio Examiner Z microscope equipped with a 20× water immersion objective. The laser ablation was performed either along a droplet diameter through the central inclusion, or along a short line adjacent to the inclusion. Fluorescence and differential interference contrast (DIC) images were acquired using a CCD camera (Cascade512B, Photometrics) to monitor the response and subsequent recovery of the F-actin network and central inclusion.

### Actin-density and flow-velocity analysis

Fluorescence intensity was used as a proxy for local actin density. The density was normalized to have a peak intensity equal to 1, and any increase or decrease in the density of sample was detected as a change in intensity. We azimuthally averaged pixel intensity in each frame to see how the intensity changes as a function of distance from the droplet center. To find the velocity of the actin filaments, we used Open PIV-matlab software with an interrogation window size equal to 16 pixels and spacing or overlap of 8 pixels (*41*). Therefore, the velocity matrix was 64 (8×8) times smaller than the original image. To reduce the noise, we ran PIV in four regions of interest (ROI) with only two pixels difference between them, and we averaged over the four velocity matrices. To see how the velocity of filaments depended on their distance from the center of the droplets, we azimuthally averaged the magnitude of velocities measured at the same radius.

### Droplet morphology and bead-tracking analysis

The image analysis was performed using Fiji/ImageJ. Droplet and inclusion boundaries were traced in equatorial optical sections. The equivalent inclusion radius *R*_1_and droplet radius *R*_2_ were calculated as 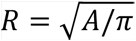, where A is the measured cross-sectional area. Fluorescent beads were tracked frame by frame to obtain trajectories and displacements. Following UV laser ablation, inclusion boundaries were fitted with ellipses, and the aspect ratio was calculated as the major -to-minor axis ratio.

## Acknowledgements

We thank Kinneret Keren, Chase Broedersz, Grzegorz Gradziuk, Margaret Cheung and Pierre Ronceray for stimulating discussions. We thank Tom Pieler, Kristine Henningfeld, Anita Smarandache and Sven Richts for help with initial *Xenopus* egg extract preparations. We thank the Light Microscopy Core Facility at Duke University, the National Xenopus Resource at the Marine Biological Laboratory in Woods Hole and the Light Microscopy Facility at the Max Planck Institute of Molecular Cell Biology and Genetics in Dresden for use of their facilities. We thank Daniel P. Kiehart for providing the device we used for magnetic tweezers experiments. We thank Dieter Klopfenstein for discussion and assistance in clarifying and documenting the *LifeAct-GFP2* purification protocol. C.D. thanks Pierre Récho for valuable early discussions regarding the numerical solution of the one-dimensional radial model.

## Author contributions

C.S. devised and directed the study and wrote the manuscript together with J.H. and with input from J.Z. and C.D. and contributions from all the authors. J.Z., A.P., C.G. and R.O. performed the experiments. U.S. purified LifeAct-GFP2. F.J. directed the modeling work, and C.D., A.S. and F.J. developed the model. C.D., A.S. and I.S. implemented the numerical solvers. J.G., R.G., A.P., B.G., M.L., R.O., S.G. and J.H. analyzed the data with input from all the authors.

## Data, code and materials availability

All data needed to evaluate the conclusions of this work are graphically presented in the paper or the supplementary materials, and can be obtained upon reasonable request from the authors.

## Supplementary Materials

Figs. S1 to S8

Movies S1 to S14

Supplementary Theory

