## Supplementary Information for "Self-centering steady-state flows emerge in confined actomyosin networks"

Jianguo Zhao *et al.*

**This PDF file includes:**

Figs. S1 to S8
Movie Legends S1 to S14
Supplementary Theory

**Other Supplementary Material for this manuscript includes:**

Movies S1 to S14

#### Supplementary figures (Fig. S1–S8)

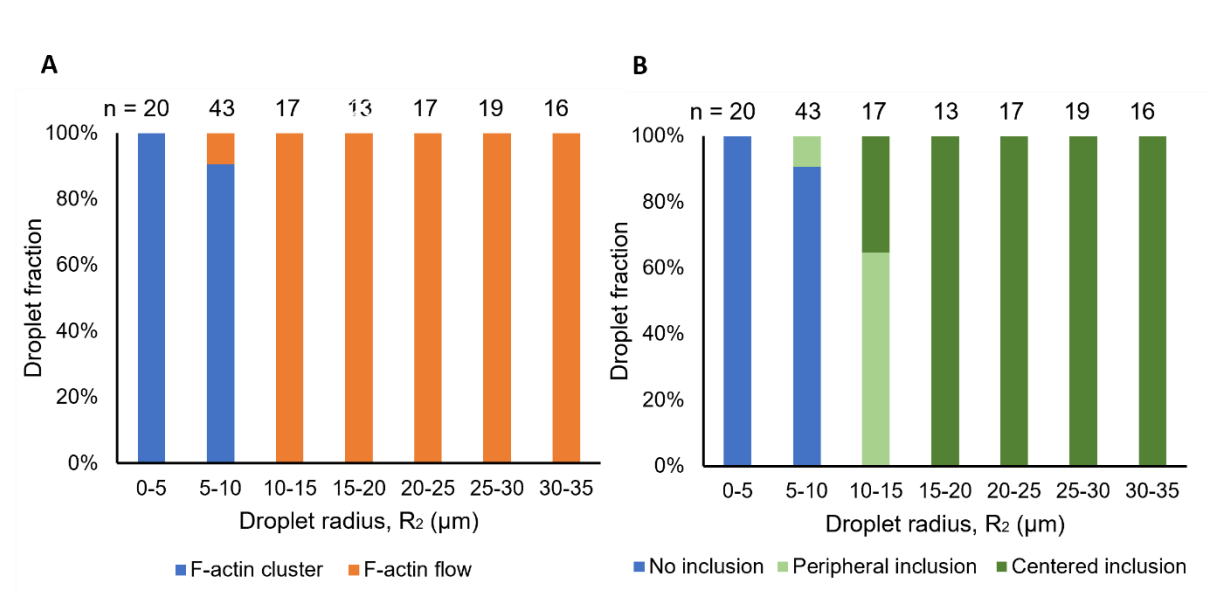

**Fig. S1. Size-dependent emergence of radially convergent F-actin flow and inclusion positioning.** (A) F-actin organization transitions from compact clustering to coherent radially convergent flow with increasing droplet size. (B) Vesicle-rich inclusions exhibit distinct positioning states as a function of droplet size. Centered inclusions occur predominantly in droplets exhibiting radially convergent flow. Data represent 145 droplets pooled from independent experiments.

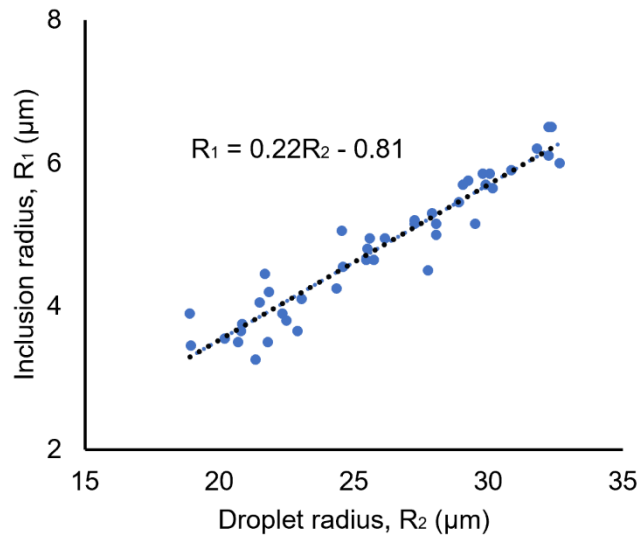

**Fig. S2. Scaling of inclusion size with droplet size.** The radius of the vesicle-rich inclusion scales approximately linearly with droplet radius, consistent with confinement-dependent organization of the steady-state F-actin flow.

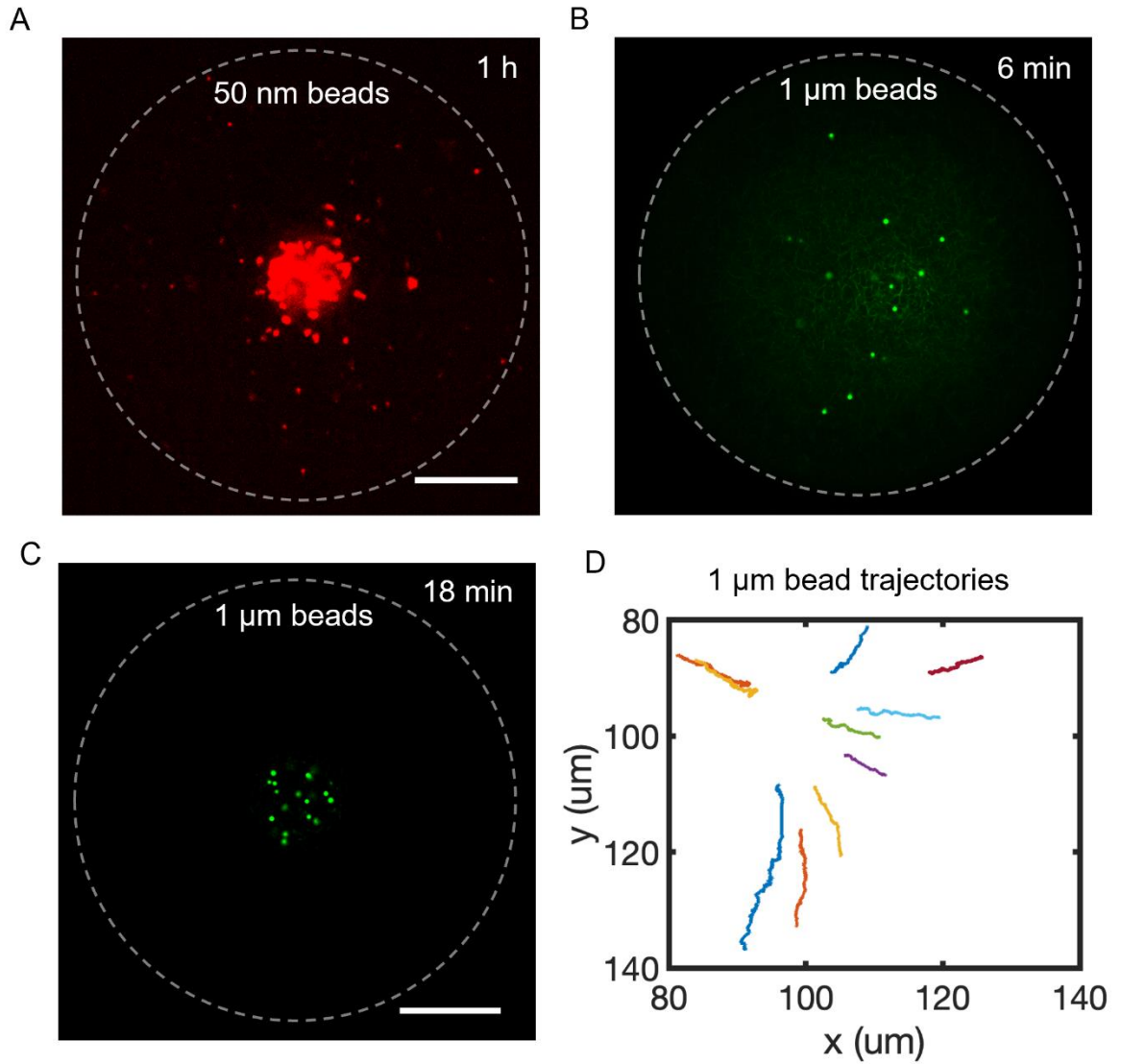

**Fig. S3. Size-dependent transport of probe beads by F-actin flow.** (A) Small probe beads (50 nm diameter) remain largely dispersed and exhibit predominantly diffusive motion 1 h after droplet formation. The Representative image was extracted from Movie S7. (B–D) Large probe beads (1 μm diameter) undergo directed transport toward the droplet center and become incorporated into the central inclusion. Representative images and trajectory data were extracted from Movie S8. Images were acquired 6 min (B) and 18 min (C) after droplet formation. Trajectories in (D) were measured over an 18 min interval. These observations are consistent with size-dependent coupling to the percolated actin network. Scale bars, 20 μm (A) and 30 μm (B, C).

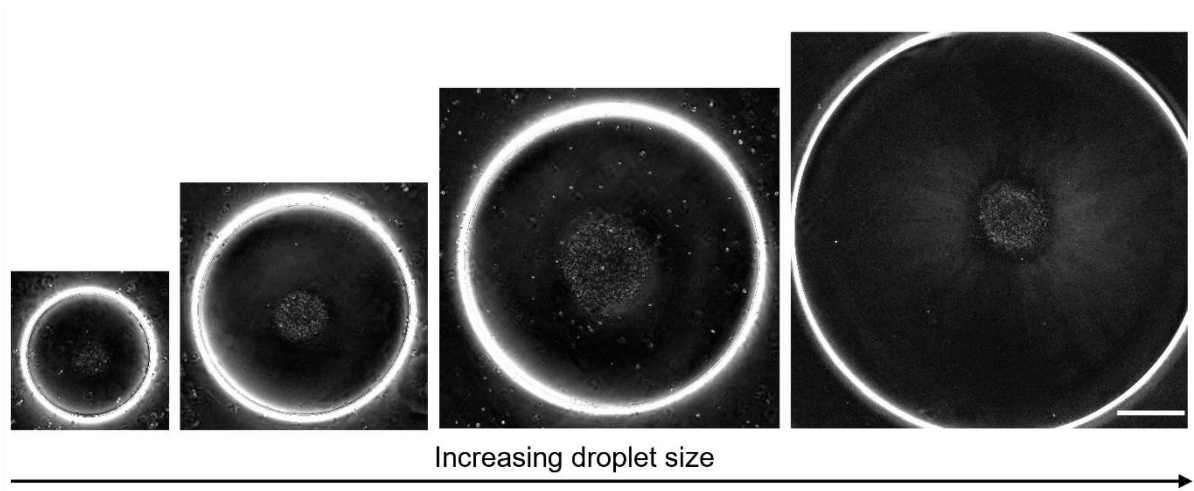

**Fig. S4. Retardance images of steady-state F-actin networks measured by LC-PolScope microscopy versus increasing droplet size.** The granular retardance pattern in the center of each droplet is caused by the lipid-rich cellular debris accumulating in the center. Outside this central pattern, no detectable birefringence was observed in the droplet interior over the droplet size range used in the quantitative analysis (left three droplets), indicating the absence of detectable large-scale nematic alignment. A retardance signal becomes measurable only in the largest droplet, where the uniform retardance decreases from 0.8 nm near the center to less than 0.1 nm near the outer edge. This centrosymmetric retardance pattern is associated with a radial slow axis alignment, indicating that the actin filaments that are expected to cause the retardance are aligned radially to the center of the droplet. Such a uniform alignment is not observed in the three smaller droplets. The strong birefringent ring observed at the water–oil interface is not accompanied by detectable F-actin enrichment in fluorescence images and arises from optical edge birefringence associated with the interface. Scale bar, 30 $\mu\text{m}$.

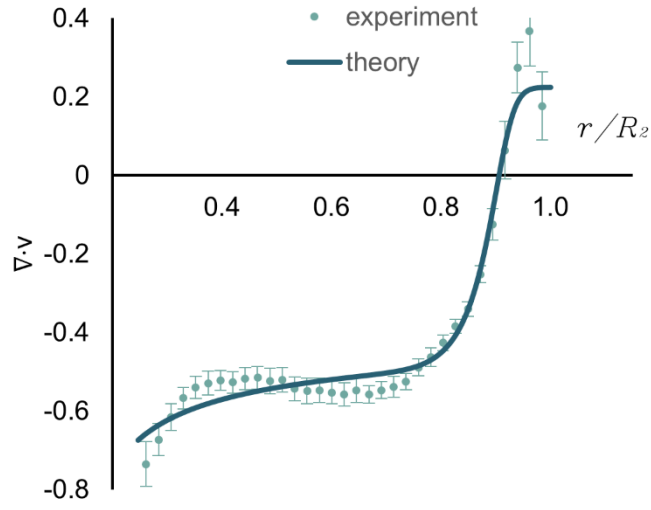

**Fig. S5. Steady-state contraction rate $\nabla \cdot v$ as a function of the normalized radial position $r/R_2$.** Experimental measurements (symbols, mean ±SD) are compared with hydrodynamic model predictions (solid line).

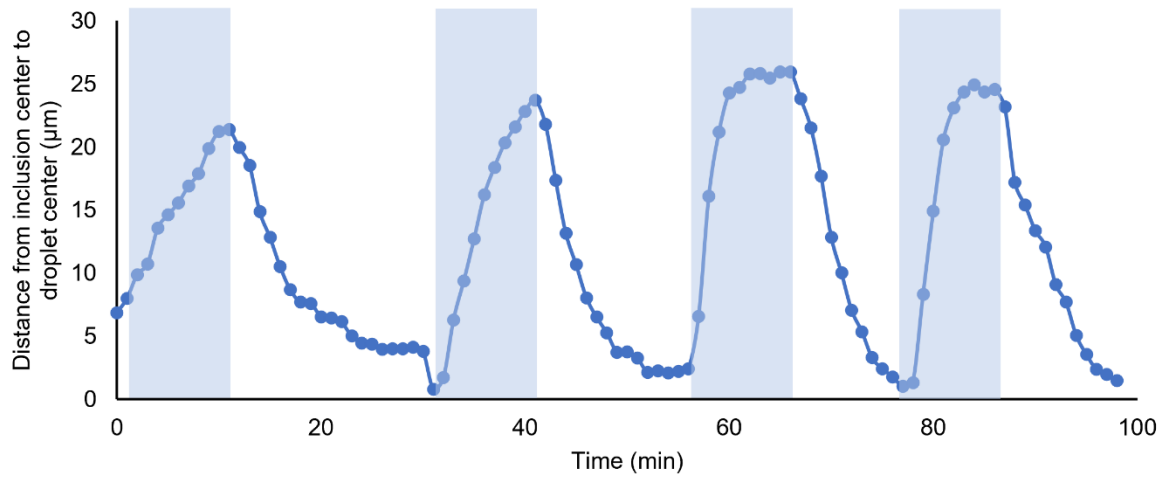

**Fig. S6. Response of the inclusion to repeated magnetic-force perturbation cycles (~300 pN).** Application of an external magnetic force displaces the inclusion away from the droplet center, whereas force removal triggers rapid recentering. The reproducible recovery dynamics demonstrate a robust self-centering steady-state maintained by the radially convergent F-actin flow. Shaded regions indicate periods of magnetic force application. Data were extracted from Movie S9.

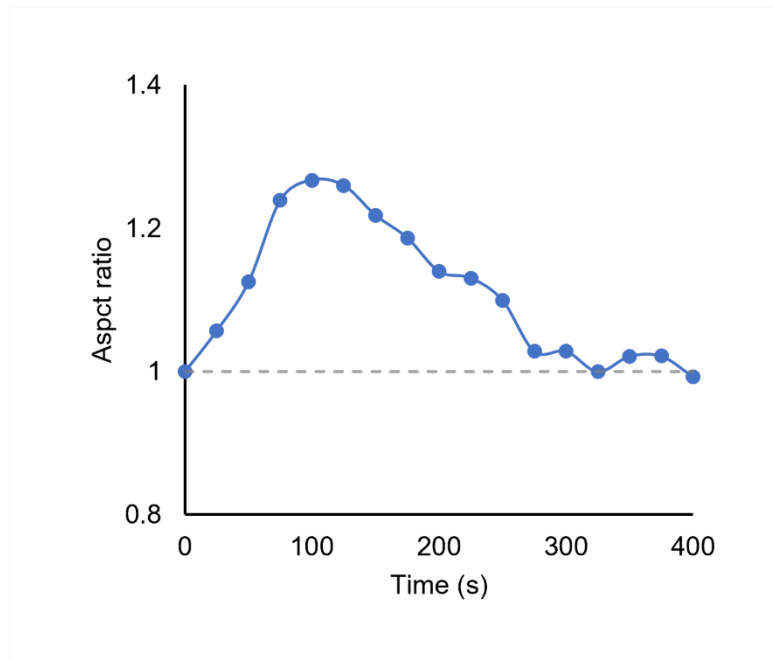

**Fig. S7. Inclusion deformation and recovery following UV ablation.** Aspect ratio of the inclusion (defined as the axis parallel to the ablation direction divided by the perpendicular axis) as a function of time following UV cutting. The transient increase and subsequent recovery of aspect ratio indicate restoration of actomyosin-generated stresses. Data were extracted from Movie S10.

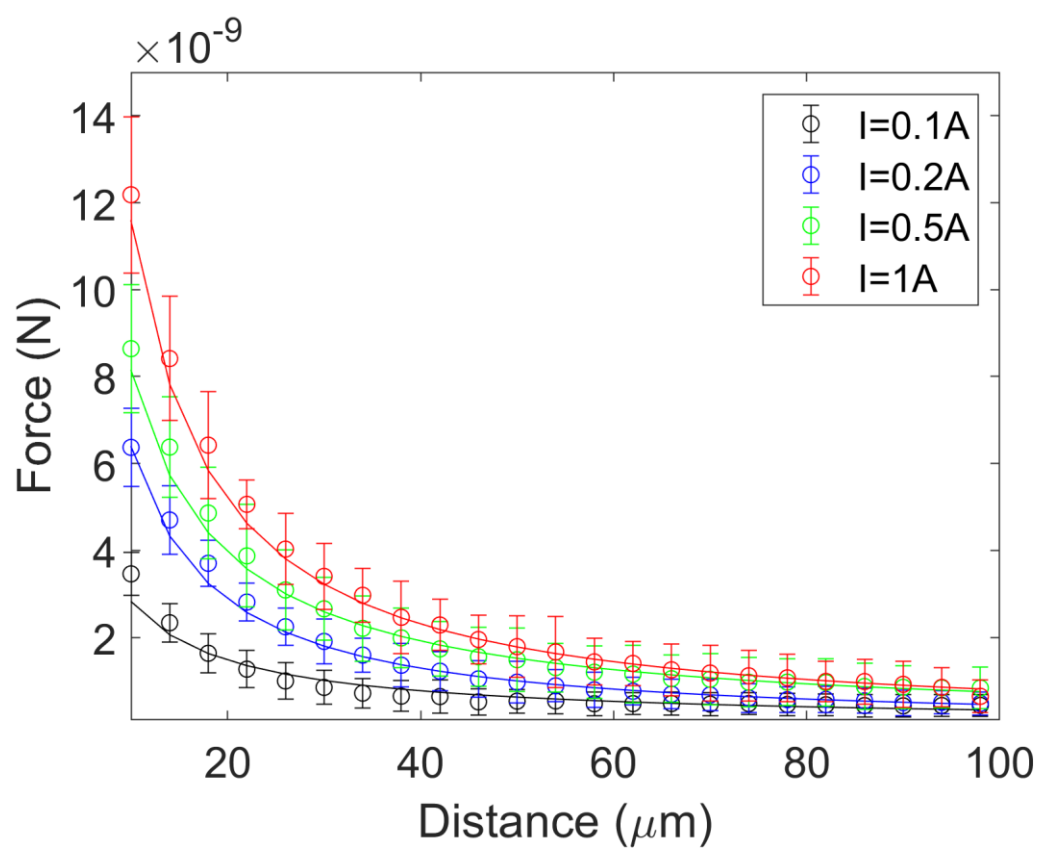

**Fig. S8. Magnetic force calibration.** Magnetic forces as a function of bead-to-tip distance at indicated solenoid currents.

#### Supplementary Movie Legends (Movie S1-S14)

**Movie S1. Small droplet exhibiting fluctuating F-actin (LifeAct-GFP, green) clusters.** Scale bar, 10 $\mu\text{m}$.

**Movie S2. Intermediate-sized droplet exhibiting peripheral inclusion positioning associated with directional F-actin (LifeAct-GFP, green) flow.** Scale bar, 10 $\mu\text{m}$.

**Movie S3. Large droplet exhibiting coherent radially convergent flow and centered inclusion formation.** Contractile F-actin flow transports vesicles toward the droplet center, leading to the growth and stabilization of central inclusion. **Left:** F-actin (LifeAct-GFP, green) flux and subsequent depletion after inclusion formation. **Right:** Vesicle transport and accumulation into a central inclusion. Scale bar, 20 $\mu\text{m}$.

**Movie S4. Three-dimensional view of steady-state F-actin flow and central inclusion.** Three-dimensional reconstruction of the contracting F-actin (LifeAct-GFP, green) network and the lipid-rich inclusion (Cy5, red). The inclusion remains centered within the droplet during steady-state F-actin flow. The red-labeled spots in the periphery are likely small non-convected vesicles. Scale bar, 30 $\mu\text{m}$.

**Movie S5. Loss of inclusion centering following ATP depletion.** Three-dimensional z-stack rendering of a droplet over time showing the displacement of inclusion from the droplet center as ATP level decreases ($\sim 3$ h after sample preparation). Scale bar, 20 $\mu\text{m}$.

**Movie S6. Three-dimensional radially convergent F-actin flow at steady state.** Side view (xz plane) of the steady-state F-actin (LifeAct-GFP, green) flow showing three-dimensional radial convergence toward the droplet center, distinct from circulatory convection. Scale bar, 15 $\mu\text{m}$.

**Movie S7. Diffusive motion of small beads within the actin network.** Fluorescent beads (50 nm diameter) remain dispersed throughout the droplet and exhibit diffusive motion without directed inward transport or central accumulation. Images were recorded 1 hour after droplet preparation. Scale bar, 15 $\mu\text{m}$.

**Movie S8. Directed inward transport of large beads by contractile F-actin flow.** Fluorescent beads (1 $\mu\text{m}$) are advected toward the droplet center and accumulate centrally over time. Images were recorded 5 minutes after preparation. Scale bar, 30 $\mu\text{m}$.

**Movie S9. Magnetic perturbation reveals reversible decentering and restoring dynamics.** Application of a magnetic force ($\sim 300$ pN) displaces the inclusion toward the droplet periphery and transiently disrupts isotropic F-actin organization. Upon removal of the force, the active F-actin (LifeAct-GFP, green) network reorganizes and the inclusion returns to the droplet center. This perturbation/recentering cycle can be repeated for over one hour, indicating a stable self-centering steady state. Scale bar, 20 $\mu\text{m}$.

**Movie S10. Numerical simulation in 3D with off-centered inclusion.** (Left) F-actin density field. (Right) F-actin velocity field (color-coded magnitude). At steady state, F-actin density profile is asymmetric and shows an F-actin accumulation at the droplet center. See SI for details on the numerical implementation.

**Movie S11. Simulations of the 3D model showing F-actin density (left) and velocity magnitude dynamics (right) starting from an initial state lacking spherical symmetry.** The system evolves toward a centrosymmetric steady-state configuration. See SI for numerical details.

**Movie S12. UV ablation reveals contractile restoring dynamics in the actin network.** Local UV ablation of the F-actin (LifeAct-GFP, green) network induces transient deformation of the central inclusion toward the ablated region. As the network reforms, the inclusion recovers its spherical shape, consistent with radially inward contractile stresses. Scale bar, 20 $\mu\text{m}$.

**Movie S13. One-sided peripheral UV ablation induces asymmetric deformation and restoring dynamics, which reveals a pushing force drives the inward F-actin flow and centering.** Localized UV ablation at one side of the inclusion boundary induces a transient deformation that recovers as the F-actin (LifeAct-GFP, green) network rebuilds, consistent with a persistent inward force maintaining the spherical inclusion shape. Scale bar, 30 $\mu\text{m}$.

**Movie S14. Numerical simulation in 3D of the F-actin dynamics after laser ablation** (simulated here by setting the F-actin density to 0 within a parallelepipedal region on both sides of the inclusion). (Left) F-actin density field. (Right) F-actin velocity field (color-coded magnitude). F-actin density dynamics shows a recovery to the steady state after the ablation. See SI for details on the numerical implementation.

### Supplementary Theory

#### I. ISOTROPIC ACTIVE GEL MODEL

We consider a spherical water-based droplet of radius $R_2$ containing a contractile acto-myosin network and surrounded by oil. Inside this droplet, the inclusion made of displaced debris is described as a rigid sphere of radius $R_1 < R_2$ (see Fig. 1B of the main text). We model the contractile acto-myosin network as a one-component active viscous fluid composed of the polymerized actin with mass density $\rho(\mathbf{r}, t)$ and velocity $\mathbf{v}(\mathbf{r}, t)$, where $\mathbf{r}$ denotes the position vector in three dimensions.

##### A. Nonlinear isotropic model with threshold

###### 1. Actin density continuity equation

Actin is advected by the flow and can be polymerized and depolymerized, such that the polymerized actin density $\rho$ obeys the following continuity equation:

$$\partial_t \rho + \nabla \cdot (\rho \mathbf{v}) = f(\rho), \quad (\text{S1})$$

where $f(\rho)$ is the actin turnover that depends on the concentration of polymerized actin and whose precise form will be discussed below.

###### 2. Force balance and constitutive equations

Force balance in the actin gel reads:

$$\nabla \cdot \boldsymbol{\sigma} = -\gamma(\rho) \mathbf{v}, \quad (\text{S2})$$

where $\boldsymbol{\sigma}$ is the actin gel stress tensor, whose constitutive equation is given below, and $\gamma(\rho)$ is a density-dependent friction coefficient. Experimental results (see main text and SI Fig. S4) suggest that the polymerized actin network is isotropic. We therefore consider the following constitutive equation for the actin gel stress:

$$\sigma_{\alpha\beta} = 2\eta(\rho)\tilde{v}_{\alpha\beta} + [\bar{\eta}(\rho)\partial_\gamma v_\gamma - P_0 + \zeta(\rho)]\delta_{\alpha\beta}, \quad (\text{S3})$$

where Greek indices denote Cartesian coordinates and where an implicit summation over repeated indices is implied. We have introduced the shear rate tensor $\tilde{v}_{\alpha\beta} = (\partial_\alpha v_\beta + \partial_\beta v_\alpha)/2 - (\partial_\gamma v_\gamma/3)\delta_{\alpha\beta}$ and a constant osmotic pressure $P_0$. We have also introduced the active isotropic stress $\zeta$ and the shear and bulk viscosities $\eta$ and $\bar{\eta}$, which may all be dependent on the local gel density $\rho$. We now discuss this density-dependent behavior.

###### 3. Nonlinear actin turnover and threshold density

As discussed in the main text, we observe from the experiments that the actin turnover is characterized by a transition between a sparse network where actin is polymerized, and a percolated, dense network at larger densities where actin depolymerizes. We capture qualitatively this behavior with a minimal number of parameters by consider the following form for the actin turnover function:

$$f(\rho) = k_p - k_d \rho - k_s \theta_w(\rho - \rho_c), \quad (\text{S4})$$

where we have introduced the nonlinear function $\theta_w(\rho) = [1 + \tanh(\rho/w)]/2$ that interpolates between the two linear turn-over regimes, and where $w$ denotes the width of the crossover. The rates $k_{p,d,s}$ are positive and denote the polymerization rate, the linear degradation rate and the stress-induced depolymerization rate. The coefficients $k_{p,d,s}$, $w$ and $\rho_c$ are obtained by the fitting the experimental data using the function *FindFit* of *Mathematica*. The fitted parameters are summarized in Table S1 and the resulting fitting curve is shown in Fig. 3A of the main text.

###### 4. Nonlinear active stress

Based on the experimental evidence of a transition between two states of the actin network, we postulate that the mechanical properties of the network also exhibit two distinct regimes separated by a nonlinear transition. Below the percolation density, the actin network is too sparse to exert active forces, and essentially behaves as a passive viscous fluid, while above it, the network is sufficiently dense to exert contractile stresses that can create fluid flows. We therefore consider the following density-dependence of the isotropic active stress:

$$\zeta(\rho) = [\mu_0 + \mu_1 \rho] \theta_w(\rho - \rho_c), \quad (\text{S5})$$

such that $\zeta(\rho) \simeq 0$ for $\rho < \rho_c$, and $\zeta(\rho) \simeq \mu_0 + \mu_1 \rho$ for $\rho > \rho_c$. In principle, the friction $\gamma$, and the shear and bulk viscosities $\eta, \bar{\eta}$ are density dependent, and they may have a similar nonlinear behavior. To limit the number of parameters in our model, we have however neglected this density dependence, and we write $\gamma(\rho) = \gamma_0$, $\eta(\rho) = \eta_0$, $\bar{\eta}(\rho) = \bar{\eta}_0$. A discussion regarding the number of parameters in our model and their estimation is given in Secs. ID, IE.

###### B. Spherically-symmetric case

In the spherically-symmetric case, the coupled equations (S1)-(S2) take a simpler form. The actin density $\rho = \rho(r)$ becomes a function of the radial distance to the centre $r = |\mathbf{r}|$, and the actin velocity $\mathbf{v} = v(r)\hat{\mathbf{r}}$ is purely radial with $\hat{\mathbf{r}} = \mathbf{r}/r$. Actin continuity equation (S1) and force balance (S2) can thus be rewritten as:

$$\partial_t \rho + \frac{2v\rho}{r} + \partial_r(\rho v) = f(\rho), \quad (\text{S6})$$

$$\partial_r \left[ \frac{2v}{r} (\bar{\eta} - 2\eta/3) + (4\eta/3 + \bar{\eta}) \partial_r v \right] + \frac{4\eta}{r} (\partial_r v - v/r) + \partial_r \zeta = \gamma v. \quad (\text{S7})$$

###### C. Boundary conditions

To solve the system of nonlinear differential equations (S1)-(S5), we need to specify three boundary conditions, which have to be imposed at the boundary with the outer droplet and at the boundary with inclusion.

###### 1. Actin concentration

The first boundary condition we impose is a fixed actin concentration at the surface $\mathcal{S}_1$ of the inclusion:

$$\rho(\mathbf{r} \in \mathcal{S}_1, t) = \rho_1. \quad (\text{S8})$$

However, since the flow resulting from actin contraction is directed inwards (towards the inclusion), imposing the boundary condition at the inclusion can be challenging. Depending on the simulation method used (see Sec. II for details), we have alternatively imposed the actin density at the outer surface $\mathcal{S}_2$, $\rho(\mathbf{r} \in \mathcal{S}_2, t) = \rho_2$.

In the spherically-symmetric case, the inclusion is centred at the origin and the boundary conditions reads:

$$\rho(r = R_1, t) = \rho_1, \quad (\text{S9})$$

or $\rho(r = R_2, t) = \rho_2$ when the boundary condition is imposed at the outer surface.

###### 2. Actin velocity

We have considered both fixed velocity and fixed stress boundary conditions. Fixed velocity boundary condition is easier to implement numerically, and the fixed values for the velocities are directly taken from the experimental data. Since the osmotic pressure $P_0$ is difficult to estimate, it can be adjusted to go from velocity boundary conditions to stress boundary conditions.

The results presented in the manuscript were thus obtained imposing two boundary conditions on the actin velocity $\mathbf{v}$, one at the surface of the inclusion and one at the outer surface of the droplet:

$$\mathbf{v}(\mathbf{r} \in \mathcal{S}_{1,2}, t) = \mathbf{v}_{1,2}, \quad (\text{S10})$$

where $\mathbf{v}_{1,2}$ are the prescribed velocity fields at the inclusion and outer surfaces. In the spherically-symmetric case, these boundary conditions simplify and read:

$$v(r = R_{1,2}, t) = v_{1,2}. \quad (\text{S11})$$

###### D. Estimation of the parameters

The shear and bulk viscosities of polymerized actin can be estimated using actin turn-over time $\tau \simeq 30 - 100$ s and the bulk modulus of the polymerized actin network $E \simeq 10^3$ Pa [1]. It yields: $\eta_0 \simeq 10^3 - 10^4$ Pa.s.

The friction coefficient $\gamma_0$ can be estimated as $\gamma_0 \simeq \eta^f/a^2$ with $\eta^f \simeq 100\eta^w$ is the viscosity of the droplet material (with $\eta^w \simeq 1$ mPa.s the viscosity of water), and $a$ is the actin meshsize. According to Ref. [2], the meshsize can be estimated using $a \simeq 1/\sqrt{\rho}$ with $\rho$ the actin concentration. We obtain: $\gamma_0 \simeq 10^{11} - 10^{13}$ kg/(m<sup>3</sup>.s).

The contractile stress $\mu_0$ can be estimated from different papers. Using Ref. [1], we can compute it using the fact that cortical tension $T \simeq 100$ pN/$\mu\text{m}$ with a cortex thickness $h \simeq 100 - 1000$ nm. It yields a contractile stress $\sigma \simeq T/h \simeq 10^2 - 10^3$ Pa. In Ref. [3], an active stress $\sigma \simeq 60$ pN/$\mu\text{m}^2 = 60$ Pa is estimated. Finally, in Ref. [4], the ratio between active contractile stress $\xi\Delta\mu$ and friction $\gamma$ is estimated as $\xi\Delta\mu/\gamma \simeq 30$ $\mu\text{m}^2/\text{s}$. Using our value of friction $\gamma \simeq 10^{11} - 10^{13}$ kg/(m<sup>3</sup>.s), we thus obtain $\xi\Delta\mu \simeq 3 - 300$ Pa.

###### E. Number of parameters and discussion

Our model involves 11 dimensionful parameters: $\gamma_0$ (friction coefficient), $\eta_0$ and $\bar{\eta}_0$ (shear and bulk viscosities), $k_p$, $k_d$ and $k_s$ (actin polymerization and depolymerization rates), $\rho_c$ (threshold density), $w$ (crossover width), $\mu_0$ and $\mu_1$ (active stress coefficients), $P_0$ (osmotic pressure). Note that the osmotic pressure $P_0$ is a constant term that does not enter force balance equations and can thus be discarded for this discussion.

Once the actin turnover function (S4) has been fitted using the steady-state mass flux divergence $\nabla \cdot (\rho\mathbf{v})$ obtained from the experiments, our model still involves 4 dimensionless parameters appearing in force balance: the dimensionless viscosity ratio $\hat{\eta} = \bar{\eta}_0/\eta_0$, the dimensionless friction $\hat{\gamma} = \gamma_0 R_0^2/\eta_0$, and the dimensionless active stress parameters $\hat{\mu}_0 = \mu_0 R_0/(V_0 \eta_0)$ and $\hat{\mu}_1 = \mu_1 R_0 \rho_0/(V_0 \eta_0)$.

Estimation of the parameters (see Sec. ID) indicate that the permeation length $L_0 = \sqrt{\eta_0/\gamma_0} \simeq 10^{-4} - 10^{-3}$ m is much larger than the system size. Friction can thus be neglected in our case but may become important for larger droplets. We therefore have set $\hat{\gamma} = 0$. Note that simulations were also performed with nonvanishing values of $\hat{\gamma} \leq 0.1$ without significant changes of the results. In addition, we expect the ratio of bulk and shear viscosities $\hat{\eta}$ to be of order 1. In the simulations, we observe that changing $\hat{\eta}$ can be compensated by changing the values of $\hat{\mu}_{0,1}$. We have therefore chosen to fix $\hat{\eta} = 1$ to avoid overfitting.

With the choice $\hat{\gamma} = 0$ and $\hat{\eta} = 1$, the values of the remaining parameters $\hat{\mu}_{0,1}$ are obtained unambiguously from fitting the density profile $\rho(r)$ and radial velocity profile $v(r)$. Table S1 summarizes the values of the dimensionless parameters obtained from the fits and the estimated values of the dimensionful ones. Finally, we emphasize that good fits cannot be obtained if either $\hat{\mu}_0$ or $\hat{\mu}_1$ is set to 0, hence highlighting the crucial role of the nonlinear transition captured by our model.

| Dimless param. | Fitted value | Comment | Dimful param. | Value | Comment |
| --- | --- | --- | --- | --- | --- |
| $\rho_c/\rho_0$ | $\simeq 0.25$ | Percolation density | $\rho_0$ | $\max(\rho(\mathbf{r}))$ | Maximum density is set to 1 |
| $w/\rho_0$ | $\simeq 0.03$ | Transition function width | $R_0$ | $\simeq 46$ $\mu\text{m}$ | Average droplet outer radius ($R_2$) |
| $k_p/f_0$ | $\simeq 0.27$ | Polymerization rate | $V_0$ | $1$ $\mu\text{m}/\text{s}$ | Reference velocity |
| $k_d/\tau_0$ | $\simeq 0.41$ | Linear depolym. rate | $\tau_0$ | $\simeq 46$ s | Reference time ($\tau_0 = R_0/V_0$) |
| $k_s/f_0$ | $\simeq 0.24$ | Stress-induced depolym. rate | $f_0$ | $\rho_0 \tau_0$ | Turnover function normalization |
| $\bar{\eta}_0/\eta_0$ | $= 1$ | Viscosity ratio | $\eta_0$ | $10^3$ Pa.s | Reference viscosity |
| $\gamma_0 R_0^2/\eta_0$ | $= 0$ | Friction coefficient | $\zeta_0$ | $\simeq 23$ Pa | Reference stress ($\zeta_0 = V_0 \eta_0/R_0$) |
| $\mu_0/\zeta_0$ | $\simeq 1.5$ | Active stress coefficient | | | |
| $\mu_1 \rho_0/\zeta_0$ | $\simeq 0.60$ | Active stress coefficient | | | |

TABLE S1. **Left.** Dimensionless parameters used in the main text and their fitted value (see Sec. IIC for details on the fitting procedure). Note that the values of $\bar{\eta}_0/\eta_0$ and $\gamma_0 R_0^2/\eta_0$ have been fixed, see Sec. IE for details. All the theory curves displayed in the main text were obtained using these values. **Right.** Dimensionful parameters and their definition.

#### II. NUMERICAL METHODS

##### A. One-dimensional solver in the spherically-symmetric case

In the spherically-symmetric case, the differential equations are effectively one-dimensional and we have solved them numerically using finite-difference schemes. We have implemented both a steady-state and a dynamical solver.

###### 1. Steady-state solver

The steady-state solver was implemented in the following way:

• the one-dimensional (dimensionless) interval $[\frac{R_1}{R_2}, 1]$ is discretized using $N$ points: $r \rightarrow \{r_1 = \frac{R_1}{R_2}, \dots, r_N = 1\}$. We typically used $N = 100$ although larger values of $N$ were used to check convergence,
• the velocity and density profiles are discretized on this interval: $v(r) \rightarrow \{v_1, \dots, v_N\}$ and $\rho(r) \rightarrow \{\rho_1, \dots, \rho_N\}$,
• the spatial derivatives are discretized using a fourth-order centered finite-difference scheme with a 5-point stencil, such that Eq. (S7) and the steady-state version of Eq. (S6) are discretized over the interval and are transformed into a nonlinear system of $2N$ equations over the variables $v_i$ and $\rho_i$,
• velocity boundary conditions (S11) and a density boundary condition (S9) are imposed.
• the nonlinear set of equations is solved using the *FindRoot* solver of *Mathematica*. The initial guess for the solver is a linear density profile $\rho(r) = a + br$ and a velocity profile given by solving analytically the differential equation (S7) for this linear density profile, for constant viscosities and friction coefficient, and for $\zeta = a' + b'\rho$.

###### 2. Dynamical solver

The dynamical solver was implemented using a finite-difference discretization of the spatial derivatives, with a first-order upwind scheme for the advective terms and an explicit time-stepping scheme. Precisely, it was implemented using *Python* in the following way:

• the one-dimensional (dimensionless) interval $[\frac{R_1}{R_2}, 1]$ is discretized using $N$ points: $r \rightarrow \{r_1 = \frac{R_1}{R_2}, \dots, r_N = 1\}$. We typically used $N = 100$ although larger values of $N$ were used to check convergence,
• time is discretized: $t \rightarrow t_n = ndt$. We typically used $dt = 10^{-3}$ although smaller values were also used to check convergence,
• the velocity and density profiles are discretized on this interval: $v(r, t) \rightarrow \{v_1(t_n), \dots, v_N(t_n)\}$ and $\rho(r, t) \rightarrow \{\rho_1(t_n), \dots, \rho_N(t_n)\}$,
• starting from a linear density profile $\rho(r, t = 0) = a + br$ as initial condition, the time stepping was performed in the following manner:
  1. the discretized version of the force balance equation (S7) forms a linear system of $N$ equations for the velocity $v_i(t_n)$ which is solved using the boundary conditions (S11).
  2. the density profile is time-stepped using the discretized version of Eq. (S6). The density boundary condition $\rho(r_N, t_n) = \rho_2$ is imposed<sup>1</sup>,
  3. time is updated to $t_{n+1}$ and the time stepping procedures continues from step 1,
• the simulation ends when the density profile is converged up to a prescribed tolerance.

We have used the dynamical solver to check that the steady-state solutions found by the steady-state solver were both stable and accessible from reasonable initial conditions.

<sup>1</sup> Using the density boundary condition at $r = R_1$ (Eq. (S9)) is unstable as the velocity is directed inwards in our system.

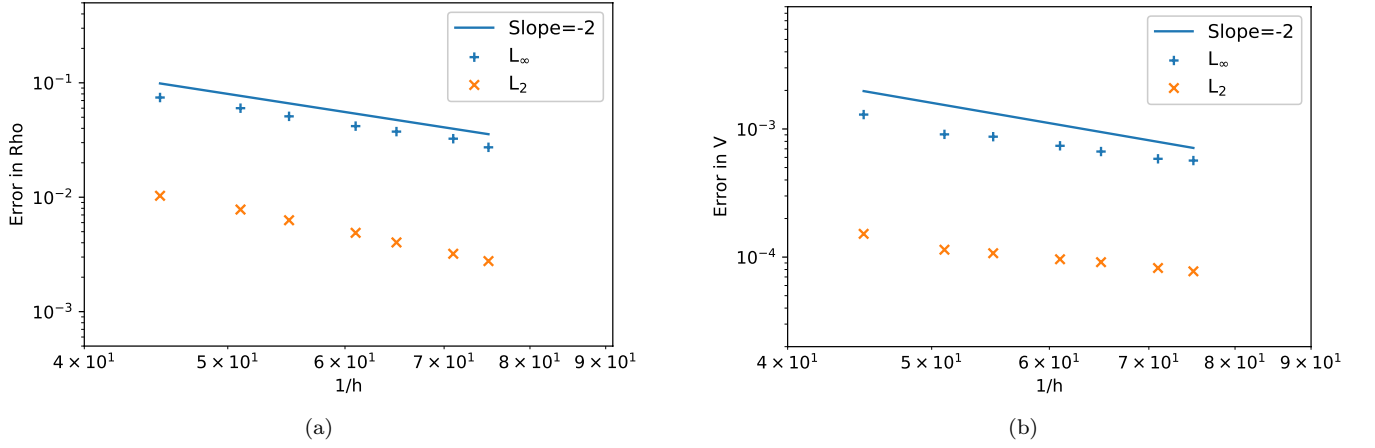

FIG. S9. Convergence of 3D dynamical solver to high-resolution numerical solution obtained from the spherically-symmetric 1D steady-state solver. (a) Convergence of the actin density. (b) Convergence of the velocity field.

##### B. Three-dimensional solver

The three-dimensional dynamical solver is implemented in the scalable computing library OpenFPM [5, 6] using the Discretization-Corrected Particle Strength Exchange method (DC-PSE) [7]. The DC-PSE method uses second order derivatives with polynomial template, with a neighbour radius $r_{\text{cut}} = 3.9h$, where $h = 2(R_2 - R_1)/(N - 1)$ with $N$ the grid size. We used $N = 55$, which correspond to an average inter-particle spacing of $r_{\text{cut}} \simeq 0.126296$, although smaller spacing were used to check convergence. The solver is implemented as follows:

• The 3D domain is discretized using Eulerian particles with uniform spacing for the domain between $(R_1, R_2)$. The spherical surface at $R_1$ is discretized by placing particles uniformly along the polar coordinates $(R_1, \theta, \phi)$, and at $R_2$ using the Fibonacci distribution technique.
• Time is discretized: $t \rightarrow t_n$ using the adaptive Adams-Bashforth-Moulten explicit predictor-corrector time-stepping method with a tolerance of $10^{-5}$.
• Starting from a radially linear initial density profile and vanishing velocity profile, time stepping was performed in the following manner:
  1. using DC-PSE for the spatial derivatives, the force balance equation (S2) is discretized and forms a linear system of equations for the velocity $v_i(t_n)$, which are solved using the Dirichlet boundary conditions (S10). On both inner and outer surface particles, each Cartesian component of the velocity is imposed through the DC-PSE linear solver.
  2. The density profile is time-stepped using Eq. (S1) and the density boundary condition $\rho(r_N, t_n) = \rho_2$ is imposed.
  3. Time is updated and time-stepping is continued.
• The simulation ends when the density profile has converged up to a prescribed tolerance.

The 3D dynamical solver has been validated against the one-dimensional steady-state solution as shown in Fig. S9.

##### C. Fitting procedure

*Actin turnover function.* The parameters that enter the definition (S4) of the sigmoid-like actin turnover have been obtained by fitting the divergence of the polymerized actin flux $\nabla \cdot (\rho \mathbf{v})$ obtained experimentally using the function *FindFit* of *Mathematica*.

*Active stress, friction and viscosities.* The four dimensionless parameters entering force balance were obtained in the following way. First, we defined a quadratic cost function that penalizes differences between the simulated density and velocity profiles and their experimental counterpart. Second, we used the *Mathematica* function *NMinimize* to find the values of the parameters that minimize this cost function. This function calls the steady-state solver described in Sec. II A 1.

Results of the fitting procedures are gathered in Table S1. All the theory curves and simulations displayed in this article have been obtained using these parameters.

###### D. Simulations without spherical symmetry

Using the 3D solver presented in Sec. II B, we can address the dynamics of a system lacking spherical symmetry. We focused in particular on the following situations:

• **Off-centered inclusion simulation.** We considered the dynamics of the F-actin density and velocity for an initially off-centered inclusion. This is displayed in Fig. 4B of the main text and Movie S13. We observe that the actin density reaches a steady state which is asymmetric and shows an enrichment of the F-actin density at the center of the droplet.
• **Laser ablation simulation.** Starting from the steady-state F-actin distribution, we simulated laser ablation by setting the F-actin density to 0 within a parallelepipedic region on both sides of the inclusion, and solved the evolution of the system from this initial condition. See main text Fig. 4D and Movie S12. Note that in our numerical model, the inclusion is a solid sphere that cannot be deformed and we do not consider the effect of the F-actin accumulation at the surface of the inclusion. The simulation, however, shows the recovery of the actin network to its spherically-symmetric steady state.
• **Heterogeneous initial condition.** We considered the dynamics of the F-actin density and velocity starting from an initial condition which lacks spherical symmetry. This is displayed in Movie S14. The dynamics drive the system to the spherical-symmetric steady state.
